# Host genetics structure Symbiodiniaceae and prokaryotic assemblages and their co-variation in sympatric fire coral species

**DOI:** 10.64898/2026.10.17.758147

**Authors:** Chloé Pozas-Schacre, Nicolas Derome, Emilie Boissin, Alexandre Mercière, Gabrielle Martineau, Pierre-Luc Mercier, Chloé A.-F. Bourmaud, Caroline E. Dubé

**Affiliations:** Institut de Biologie Intégrative et des Systèmes (IBIS), Université Laval, Québec, Canada; Université de la Polynésie française, IFREMER-LM-IRD-EIO, UMR 241 SECOPOL, Polynésie Française; PSL Université Paris: EPHE-UPVD-CNRS, UAR 3278 CRIOBE, Université de Perpignan, Perpignan, France; Laboratoire d’Excellence CORAIL, France; Université de La Réunion, UR-IRD-CNRS-UNC-IFREMER, UMR 9220 ENTROPIE, La Réunion

**Keywords:** *Millepora*, Host genotype, *Symbiodinium*, Bacterial assemblages, Microbiome integration, Symbiont co-variation

## Abstract

Host genetic and environmental filtering are key determinants of microbiome assembly, yet their relative influence across multiple symbiotic compartments remains poorly understood. In coral holobionts, studies typically focus on either algal or bacterial symbionts and rarely account for host genetic variation below the species level. Here, we combined host microsatellite genotyping with ITS2 and 16S rRNA metabarcoding to investigate the drivers of Symbiodiniaceae and prokaryotic assemblages in three sympatric fire coral species (*Millepora* cf. *exaesa*, *M. tenera*, and *M.* cf. *platyphylla*) across three adjacent but environmentally contrasting reef habitats on Réunion Island. Both symbiont assemblages were structured by host identity and habitat, although host genetics consistently outweighed environmental effects on the microbiome composition. At the subspecies level, host genotypes explained up to 54% of the variation in Symbiodiniaceae assemblages compared to 20% for the prokaryotes. Symbiodiniaceae communities were dominated by *Symbiodinium* A16 and A3 radiations, while showing marked specificity across species and habitats. All *Millepora* species consistently associated with the Spirochaetaceae, Kiloniellaceae and Desulfobacteraceae bacterial families across all habitats, suggesting evolutionarily maintained core associations. Integrative analysis further demonstrated genotype-specific co-variation between algal and bacterial symbionts, with some genotypes maintaining stable and co-varying symbionts across clones, while others exhibited more flexible assemblages. Together, our results indicate that sympatric fire coral acquire symbionts from locally available pools through host genotype-dependent filtering. This study highlights host genotype as a major determinant of coral holobiont assembly, shaping the composition and co-variation of both Symbiodiniaceae and prokaryotic assemblages.

## INTRODUCTION

Understanding the processes that govern holobiont assembly and maintain complex symbiotic communities is a central question in microbial ecology. In host-associated microbiomes, community composition emerges from the interplay between host compatibility, environmental selection and microbial interactions [1–3]. Reef-building corals provide a powerful model for investigating these processes due to their associations with diverse symbionts, including dinoflagellates (Symbiodiniaceae family) [4], bacteria, archaea, viruses and fungi [5, 6], that contribute directly to host functioning [7–9]. The Symbiodiniaceae fulfil much of the nutrition requirement of their hosts via photosynthesis [8, 10] and largely took part in the ecological success of modern coral reefs [11]. Prokaryotic symbionts contribute a wide array of functions, from nutrient cycling to metabolite provision (e.g., amino acids, vitamins), and play central roles in holobiont immunity and stress resilience [8, 9, 12]. Yet, a major unresolved question is whether host genetic and environmental filtering simultaneously act across multiple symbiotic compartments, thereby resulting in coordinated patterns of symbiont association within the coral holobiont.

Coral hosts exert strong selective pressure on their symbionts, often leading to phylosymbiotic signals of both Symbiodiniaceae [13, 14] and prokaryotic assemblages [6, 15, 16]. Such host-symbiont congruence suggests that specific symbionts provide consistent benefits to their hosts maintaining their long-term association across generations [17, 18]. Accordingly, symbiont assemblages often show high specificity to host species [2, 19–21]. However, most studies considered host genetic identity at the species level, potentially masking biologically meaningful variation among host genotypes [1, 22, 23]. Thus, whether host genetic effects remain detectable below the species level and across multiple symbiotic compartments remains poorly understood.

Environmental constraints are also strong determinants of symbiont assemblages (e.g., temperature, light, turbidity and bathymetry [1, 24–26]). Such microbiome plasticity was identified as a key mechanism for holobiont acclimatization through the acquisition of locally adaptative symbionts. While Symbiodiniaceae tend to remain host-specific [20, 21, 23, 27, 28], prokaryotic assemblages are often more plastic, albeit host-specific patterns [1, 21–23]. In fire corals, genetically identical colonies (clones) exhibited plastic bacterial assemblages across reef habitats [1]. Yet, host genetics can modulate the extent of this environmental responsiveness [2, 23, 29], highlighting the need for integrative approaches that simultaneously consider intraspecific host genetic variation and multiple symbiotic compartments to understand coral holobiont assembly.

Despite increasing recognition that coral fitness emerges from interactions among multiple microbial partners [30], most studies have examined Symbiodiniaceae and prokaryotic assemblages independently, leaving their interactions largely unexplored [30]. While little correlation between these two symbiotic compartments has been reported [25, 31], recent work suggests that different Symbiodiniaceae lineages are associated to distinct bacterial assemblages [3, 32]. Yet, it remains unclear whether host genetic variation coordinately structures both symbiotic compartments within coral species. Naturally occurring fire coral clones across contrasting environments [1, 27, 33] provide a rare opportunity to disentangle the interplay of host genetic and environmental filtering on host-symbiont associations, a challenge that is difficult to address in most coral systems.

Here, we used ITS2 and 16S rRNA metabarcoding with microsatellite genotyping of the host to characterize the structure of prokaryotic and Symbiodiniaceae assemblages in three *Millepora* species on Réunion Island: *Millepora cf. exaesa*, *M. tenera*, and *M. cf. platyphylla*. These sympatric species are distributed across three adjacent, but environmentally distinct reef habitats: the inner and outer reef flats and the mid reef slope. This framework allowed us to examine how host–symbiont associations are shaped across genetically distinct hosts and natural environmental gradients. We first examined the contribution of host genetics (i.e., species and genotypes) and the environment on the diversity and composition of each symbiotic assemblage. At the genotype level, we investigated whether both symbiotic compartments responded in coordinated ways by looking at patterns of co-dispersion and co-variation.

## MATERIALS AND METHODS

### Experimental design and coral sampling

Between April 13 and May 24 2017, 192 colonies of *Millepora* cf. *exaesa*, *Millepora tenera* and *Millepora* cf. *platyphylla* (hereafter referred to as *M. exaesa*, *M. tenera* and *M. platyphylla* for simplicity) were mapped and sampled from the inner reef flat (1-2 m depth; IRF), outer reef flat (< 1m depth; ORF) and mid reef slope (10-15m depth; MRS) at the site “Kiosque” on the west coast of Reunion Island, France (St-Leu; 21° 10’56’’S 55° 17’08’’E; Fig. 1; Table S1). Species boundaries were assessed morphologically in the field and further verified by sequencing of the mitochondrial 16S gene (Fig. S1)

**Figure 1.**
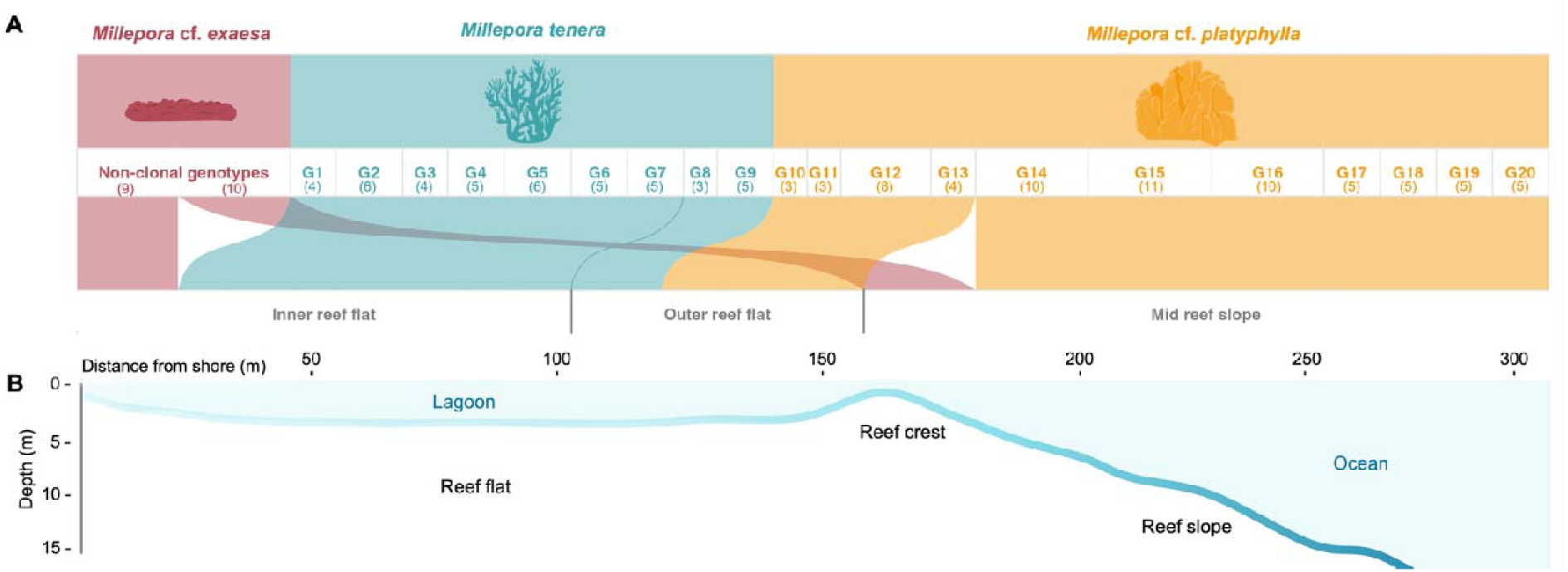
Distribution of *Millepora* spp. colonies belonging to three species and across three reef habitats. **(A)** Colonies of *M.* cf. *exaesa, M. tenera* and *M.* cf. *platyphylla* were mapped and sampled in three adjacent habitats: the inner reef flat (1-2 m depth; IRF), outer reef flat (< 1 m depth; ORF) and mid reef slope (10-15 m depth; MRS). The thickness of the ribbons corresponds to the number of replicates per species or genotype in the habitat in which they were found. Number of replicates for non-clonal and clonal genotypes (G1 to G20) are indicated in parenthesis. For *M.* cf. *exaesa*, genotypes were non-clonal in contrast to the *M. tenera* and *M.* cf. *platyphylla*. **(B)** Illustrative depth (m) and distance from shore (m) profiles of the study site “Kiosque” at St-Leu, on the west coast of Reunion Island (21° 10’56’’S 55° 17’08’’E).

Within each habitat, three 100 m long and 10 m wide belt transects were laid parallel to the shore. Each colony was mapped according to its x and y coordinates representing its position along the transect width (0-10 m) and length (0-100 m), respectively. Small fragments (< 2 cm^3^) were chiseled from each colony and placed into 2 mL tubes underwater. Each fragment was rinsed with 70% ethanol to remove bacterioplankton and preserved into 80% ethanol until DNA extraction.

### DNA extraction, species delineation and genotype identification

Fragments were incubated at 55 °C for 1h in 450 µL of lysis buffer and proteinase K (QIAGEN, Germany). Genomic DNA was then extracted using a QIAxtractor automated-extraction instrument according manufacturer’s instructions.

Species identity was verified by sequencing the mitochondrial 16S gene using the 16S-SHA and 16S-SHB primer, following Boissin *et al.* (2022) [33], who provided morphological and molecular delineation of *M. exaesa*, *M. tenera*, and *M. platyphylla* from Réunion Island. PCR reactions (25 µL), containing 2 µL of template DNA, were performed on all *M. exaesa* samples (*n* = 19) and one representative sample per clonal genotype for *M. tenera* (*n* = 11) and *M. platyphylla* (*n* = 11). For *M. tenera*, one additional clone of G4 and G6 were sequenced due to variability in the dominant Symbiodiniaceae within these genotypes (see Results; Fig. 2B). The PCR mix composition and thermal cycling conditions used are described in Ory *et al.* 2025 (detailed in the Supplementary Methods) [34].

**Figure 2.**
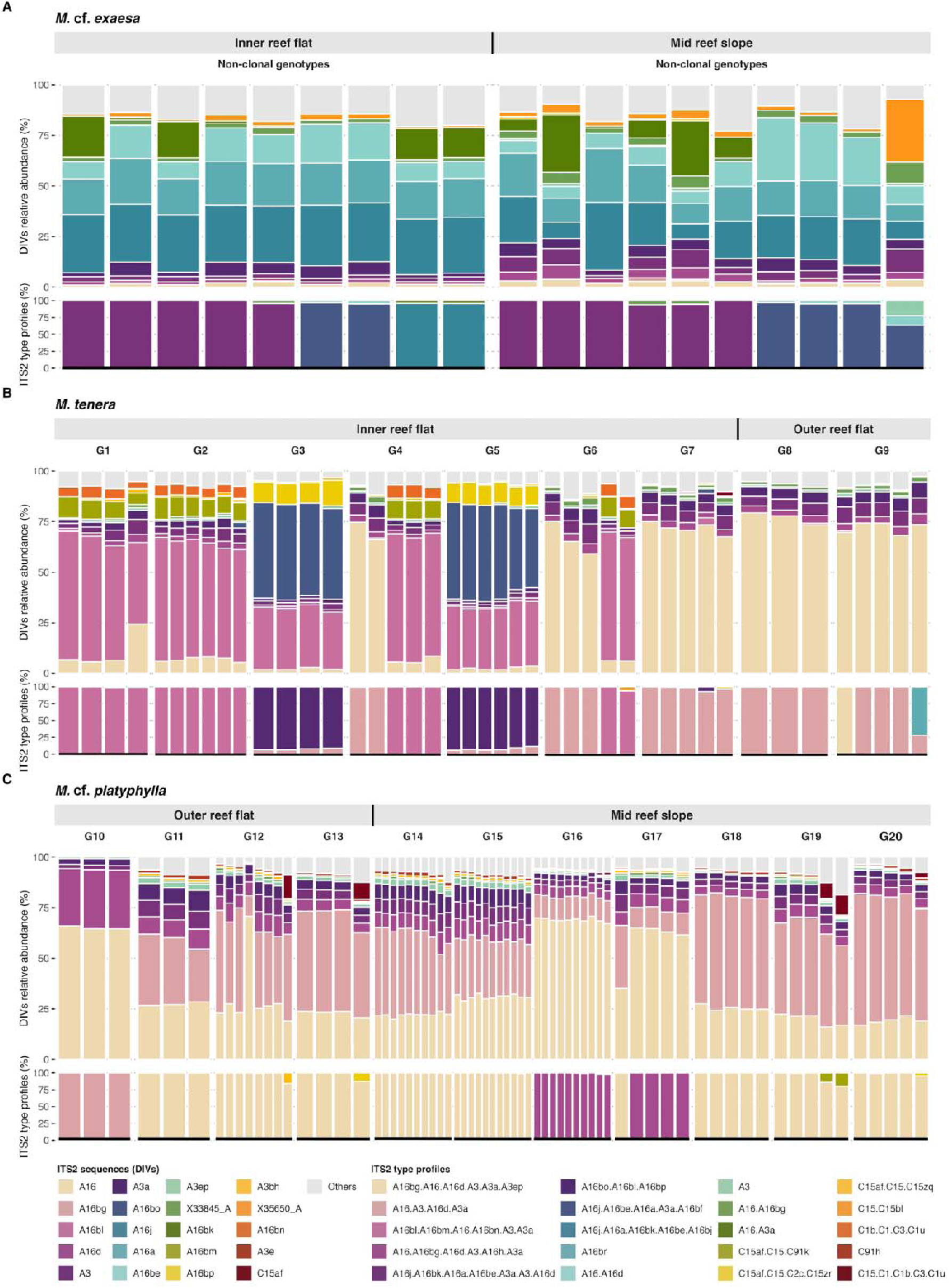
Composition of Symbiodiniaceae assemblages in *Millepora* spp. across host species, genotypes and reef habitats. Each panel corresponds to the relative abundance of post-MED ITS2 sequences (DIVs; top barplots) and of predicted ITS2 type profiles (bottom barplots) in each sample of the three species **(A)** *M.* cf. *exaesa*, **(B)** *M. tenera* and **(C)** *M.* cf. *platyphylla.* Genotypes are grouped by habitats. The 20 ITS2 sequences and 20 type profiles with the highest mean relative abundance are shown in the legends, with other sequences and profiles grouped in the “Other” category.

Briefly, the phylogenetic tree was inferred based on Maximum likelihood using IQ-TREE2 and the best-fit substitution model identified by ModelFinder [35, 36]. Species assignation was verified by reference sequences from Reunion Island available on GenBank [33]. A sequence of *Apatizanclea divergens* (Hydrozoa, Zancleidae family) was used as an outgroup to root the phylogenetic tree (see Fig. S1 and Data availability section for GenBank accession numbers).

Multilocus genotypes (MLGs) were identified from the amplification of 9, 15 and 17 polymorphic microsatellite loci for *M. exaesa*, *M. tenera* and *M. platyphylla*, respectively [37, 38]. MLGs detected in a single colony were considered unique. Clonal genotypes were observed only for *M. tenera* and *M. platyphylla*, for which we selected only those with at least 3 clones for further analysis (see Fig. S2 and Supplementary Methods for details). Raw microsatellite data is available in Table S2.

### ITS2 and 16S rRNA genes amplification and sequencing

Symbiodiniaceae assemblages were characterized by amplifying the Internal Transcribed Spacer 2 (ITS2) using the forward SYM_VAR_5.8S2 and reverse SYM_VAR_REV primers [39]). Prokaryotic assemblages were characterized by amplifying the V4 regions of the 16S rRNA gene using the primers 519F and 803R. PCR reactions of 50 µL were prepared containing 3 µL of DNA template and 2.5 µL of each primer (10 µM) using the Q5 High-Fidelity PCR kit (New England Bio Labs). PCR products were paired-end sequenced (2 x 250bp) using Illumina NovaSeq 6000 technology at CHUL Québec, Canada. See the Supplementary Methods for primer sequences and PCR conditions.

### Metabarcoding data processing

Raw ITS2 sequencing reads were submitted to SymPortal for quality control and analysis [51]. The pipeline produced defining intragenomic variants (DIVs) and inferred type profiles (i.e., proxy for Symbiodiniaceae genotypes) for each colony [40]. We filtered out singleton sequences, resulting in a dataset with an average of 494,179 reads per sample across 245 DIVs (Table S1), which can be found in Table S3.

DADA2 was used for quality filtering, denoising, merging of 16S rRNA gene reads, inference of amplicon sequence variants (ASVs) and chimera removal [41]. Briefly, forward and reverse reads were truncated at 249 bp each, quality filtered with a maximum expected error of 2 and a quality score threshold of 2. Taxonomic assignation of ASVs was conducted at the genus-level against the *Silva* v.138.1 database. Eukaryotic ASVs (i.e., chloroplasts or mitochondria), putative contaminants (decontam R package [42]) and singletons were removed prior to analysis. We further filtered out the data by removing samples with less than 10,000 reads, resulting in *n* = 131 samples, including 19 *M. exaesa,* 43 *M. tenera* and 69 *M*. *platyphylla*. This yielded 450,819 reads per sample on average across 43,387 ASVs in total, belonging to 327 families and 627 genera (Table S1). The phyloseq R package was used to combine the taxonomic table, the ASV abundance data and the metadata [43]. ASV raw abundances across each sample and taxonomic information can be found in Table S4.

### Symbiodiniaceae and prokaryotic community analysis

The Symbiodiniaceae community was characterized from the post-MED abundance data. We generated stacked barplots representing the mean relative abundance of the 20 most abundant ITS2 DIVs and profile types. For the prokaryotic community structure, we first examined the core microbiome using a 80% occurrence threshold [44]. A core was identified within each host species-habitat combination (i.e., total of 6) to highlight key bacterial symbionts in a given environmental niche. Core ASVs were taxonomically grouped by summing the abundances of individual ASVs at the lowest taxonomic level. Additionally, we generated stacked barplots representing the mean relative abundance of the 20 most abundant prokaryotic families.

We tested for differences in alpha-diversity (Shannon diversity) on count data rarefied at the minimum library size. At the species-level, we used linear models for prokaryotic assemblages, with species, habitat and their interaction as fixed factors, and Kruskal-Wallis tests for Symbiodiniaceae assemblages due to heteroscedasticity among species. For Kruskal-Wallis, we tested separately the effects of species, habitats and their combination (i.e., combined species-habitat variable), followed by Dunn post-hoc tests with Benjamin-Hochberg (BH) correction. At the subspecies-level, because host genotypes were habitat-specific, we used nested linear models to test the effect of habitat and host genotype nested into habitats, separately for *M. tenera* and *M. platyphylla*. Assumptions of Gaussian distribution and homoscedasticity were verified with the DHARMa R package [45]. Significance of fixed effects was assessed by ANOVA and posthoc comparisons by Tukey’s HSD (emmeans R package; [46]).

For multivariate analysis, DIV and ASV raw abundances were transformed using the centered-log ratio (CLR) to account for the compositional nature of the metabarcoding data (microbiome R package; [47]). Data was transformed on subsetted datasets and after the removal of taxa that summed to zero. We investigated the influence of host genetics, at both the species- and subspecies-level (i.e., genotypes) and reef habitats on the composition of Symbiodiniaceae and prokaryotic assemblages. At the species-level, we conducted a two-way PERMANOVA with the fixed factors species, habitats and their interaction (999 permutations; vegan R package; [48]). Due to the unbalanced design, significance of each factor was assessed using marginal effects (by = “margin”), ensuring that the order of factors in the model did not influence the results. Pairwise PERMANOVA were subsequently used to identify differences between group levels. At the subspecies-level, we applied a nested multivariate model with habitats and genotypes nested into habitats as fixed factors (999 permutations). To assess whether significant PERMANOVAs were due to heterogeneous dispersions, we tested for differences in group dispersions with “betadisper” and permutation tests (999 permutations). Microbial community structure was visualized using principal component analysis (PCA) from Euclidean distances of CLR-transformed abundances.

To identify symbionts that contributed most to species and habitats discriminations, we performed sPLS-DA analysis within 1) each species for habitat discrimination and 2) each habitat for species discrimination, resulting in six supervised models. sPLS-DA were run on CLR-transformed and scaled data using two components, with the optimal number of variables selected by M-fold cross-validation (mixOmics R package; [49]. Model performance was assessed by cross-model validation and the significance of the classification error rate (CER) was tested by permutation tests (999 permutations, RVAideMemoire R package; [50]). The top10 variables with the highest VIP scores were visualized in biplots with clustered samples.

### Associations between Symbiodiniaceae DIVs and prokaryotic ASVs

Associations between Symbiodiniaceae DIVs and prokaryotic ASVs were assessed for *M. tenera* and *M. platyphylla*, for which host clonal genotypes were available. We first estimated the correlation between Symbiodiniaceae and prokaryotic dispersions at the host genotype level (Pearson’s correlation) assessing the extent to which dispersion of symbiont compartments is coordinated across clones. Within-genotype dispersions were retrieved from Euclidean matrices subsetted by species. To account for species-specific multivariate scales, dispersions values were scaled within each species. We fitted a linear model to test the influence of Symbiodiniaceae dispersion, host species and their interaction on the prokaryotic dispersion.

We then examined co-variations of Symbiodiniaceae and prokaryotic assemblages across genotypes by using the DIABLO supervised algorithm (multi-block PLS-DA, mixOmics R package [49]). DIABLO models were run separately for each species-habitat combination to identify pairs of DIVs and ASVs that both discriminated host genotypes and were correlated across datasets. We set a design matrix with a correlation of 0.1 (weight) between data blocks to prioritize discrimination. The optimal number of components and variables were tuned by cross-validation (function mixOmics::tune.block.splsda). DIABLO outputs were validated by a permutation test based on cross-model validation function (function RVAideMemoire::DIABLO.test). Key contributing DIVs and ASVs were identified by selecting those with the highest importance based on the maximum absolute loading across components. CLR-transformed abundances of selected variables were visualized using heatmaps with Euclidean distance and complete linkage clustering. Finally, we quantified sample-level concordance between Symbiodiniaceae and prokaryotic communities by computing Euclidean distances between paired sample latent scores across genotypes, reflecting how similarly each sample was positioned in the integrative space.

## RESULTS

### Contribution of host genetics and habitats on Symbiodiniaceae and prokaryotic assemblages

Symbiodiniaceae alpha diversity (Shannon index) was influenced by host species, habitats and their combination (all three Kruskal-Wallis, p < 0.05; Fig. S3A, Table S5). *M. exaesa* harbored the highest diversity across habitats, and *M. platyphylla* exhibited greater diversity than *M. tenera* in the outer reef flat. Alpha-diversity was influenced by habitat in *M. tenera* only, while host genotype within habitat was significant for both *M. tenera* and *M. platyphylla* (nested LM; p < 0.001). In contrast, prokaryotic alpha-diversity was largely consistent across host species and habitats (LM, p > 0.05; Fig. S3B; Table S5), except for *M. platyphylla* which harbored more diverse assemblages in the mid reef slope than the outer reef flat. No significant effect of host genotype was detected on alpha-diversity of prokaryotic assemblages.

Symbiodiniaceae assemblages (i.e., ITS2 DIVs) were primarily structured by host species (R^2^ = 0.14), followed by habitat (R^2^ = 0.03) and their interaction (R^2^ = 0.01; PERMANOVA; all p < 0.01; Fig. S4A: Table S6). Similarly, prokaryotic assemblages were shaped by host species (R^2^ = 0.04), habitat (R^2^ = 0.03) and their interaction (R^2^ = 0.01; PERMANOVA, p < 0.05; Fig S4B; Table S6). Each coral species harboured a distinct microbiome in each habitat. No confounding effects of multivariate dispersion were detected for either symbiotic compartments (Table S7).

At the subspecies level, host genotype far exceeded the influence of habitat on both Symbiodiniaceae and prokaryotic assemblages. Host genotype accounted for 46% and 54% of the total variation in Symbiodiniaceae assemblages of *M. tenera* and *M. platyphylla*, respectively, compared to only 3-11% for habitat (PERMANOVA, p < 0.001; Fig. S5; Table S8). For prokaryotic assemblages, host genotype explained 20% and 18% of the total variation, whereas habitat accounted for only 3-5% (Fig. S6; Table S9). Differences in prokaryotic assemblages among host genotypes were detected across all habitats, except for *M. tenera* in the outer reef flat. Although host genotypes showed heterogeneous multivariate dispersions, PCA ordinations suggest that significance in PERMANOVA emerged from compositional differences (Table S10).

### Symbiont community composition across host species

The Symbiodiniaceae community was dominated by the genus *Symbiodinium*, representing 99.2% of total reads (*n* = 109 DIVs), while *Cladocopium* accounted for only 0.8% despite a greater number of DIVs (*n* =132 DIVs). Five core *Symbiodinium* sequences, A16, A16d, A3, A3a, and A3ep, were detected in > 80% of all samples (Fig. 2). SymPortal inferred 29 type profiles (i.e., unique combinations of intragenomic sequence variants) from 60 DIVs, with *M. platyphylla* hosting 14 profiles, followed by 12 in *M. tenera* and 8 in *M. exaesa.* Despite this apparent generalism (i.e., host or symbiont that associate with a variety of partners), individual colonies typically hosted only one or two dominant profiles, reflecting a high degree of host-symbiont specificity (Fig. 2). *M. platyphylla* displayed the highest diversity and relative abundance of *Cladocopium* sequences and was the only species with rare *Fugacium* and *Gerakladium* sequences (Figs 2 & 3C; Table S3). It was dominated by a single profile (A16.bg.A16.A16d.A3.A3a.A3ep), shared by 8 of 11 genotypes, while *M. tenera* harbored three dominant profiles largely absent from *M*. *platyphylla* (except the A16.A3.A16d.A3a in G10). In contrast, *M. exaesa*, harbored entirely different profiles (e.g., A16j & A16a-based profiles) that did not overlap with the other two species, aligning with its greater phylogenetic divergence (Fig. S1). Notably, *M. tenera* genotypes that share the same dominant ITS2 profiles (e.g., A16bo.A16bl.A16bp shared by G3 and G5), also clustered together in host phylogeny (Fig. 2; Fig. S1), suggesting some host cryptic lineages associated with specific photosymbionts. Consistent with these host-specific associations, sPLS-DA clearly discriminated Symbiodiniaceae assemblages according to host species (p < 0.001; Fig. 3A-C), with A16a and A16j driving the separation of *M. exaesa* from the two other species (Figs 2 & 3A, C).

**Figure 3.**
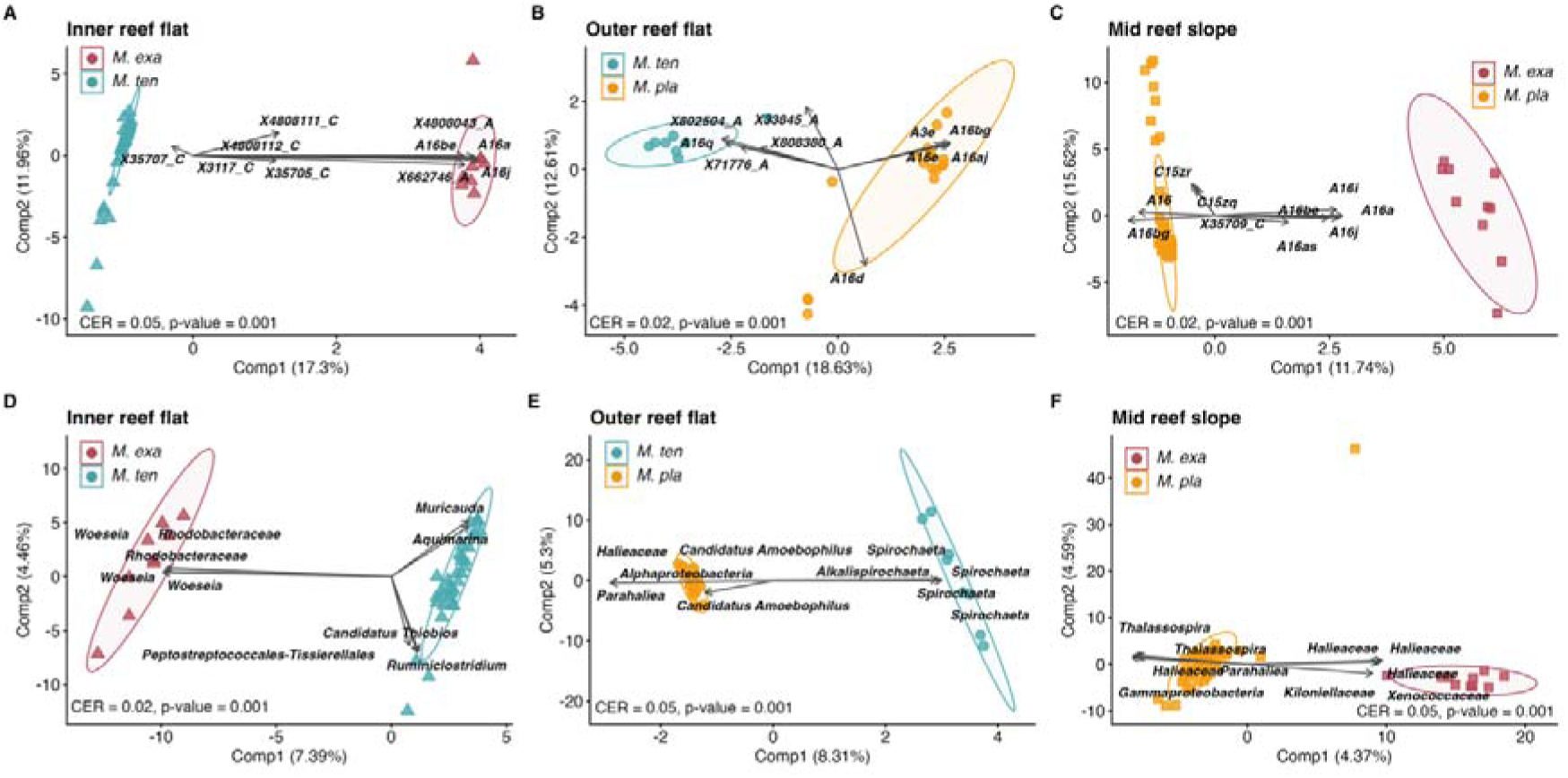
Host species discrimination based on Symbiodiniaceae and prokaryotic assemblages. sPLS-DA biplots of **(A-C)** ITS2 sequences (DIVs) and **(D-E)** ASVs illustrating the discrimination of species within each habitat. Points represent individual samples, while arrows indicate the top 5 contributing variables for each species discrimination. Count data were CLR-transformed and scaled. Classification error rate (CER) and pvalue were determined by cross-model validation and permutation tests (nperm = 999). Ellipses represent the spread of the data points at a 95% confidence level.

Prokaryotic assemblages were dominated by the Spirochaetaceae (19.6 % of total reads), Flavobacteriaceae (4.7%) and Kiloniellaceae (4.6 %) families (Fig. 4). The core microbiome consisted of 668 ASVs, including members of the *Spirochaeta* and *Alkalispirochaeta, Muricauda, Fodinicurvata* and *Tistlia,* that were detected across all host-habitat combinations (Fig. S7). The Spirochaetaceae were particularly abundant in *M. platyphylla* (e.g., up to 34.3% in the outer reef flat) and in *M. tenera* (∼17% in both reef flats), whereas Halieaceae and Tenderiaceae were particularly abundant in *M. exaesa*. Host species strongly discriminated prokaryotic assemblages in each habitat (p < 0.001; Fig. 3D-F). In the inner reef flat, *M. exaesa* was enriched in several *Woesia*, while *M. tenera* was characterized by *Aquimarina* and *Muricauda*. *M. platyphylla* was discriminated from *M. tenera* and *M. exaesa* by an enrichment of specific ASVs belonging to Haliaceae (e.g., *Parahaliea*).

**Figure 4.**
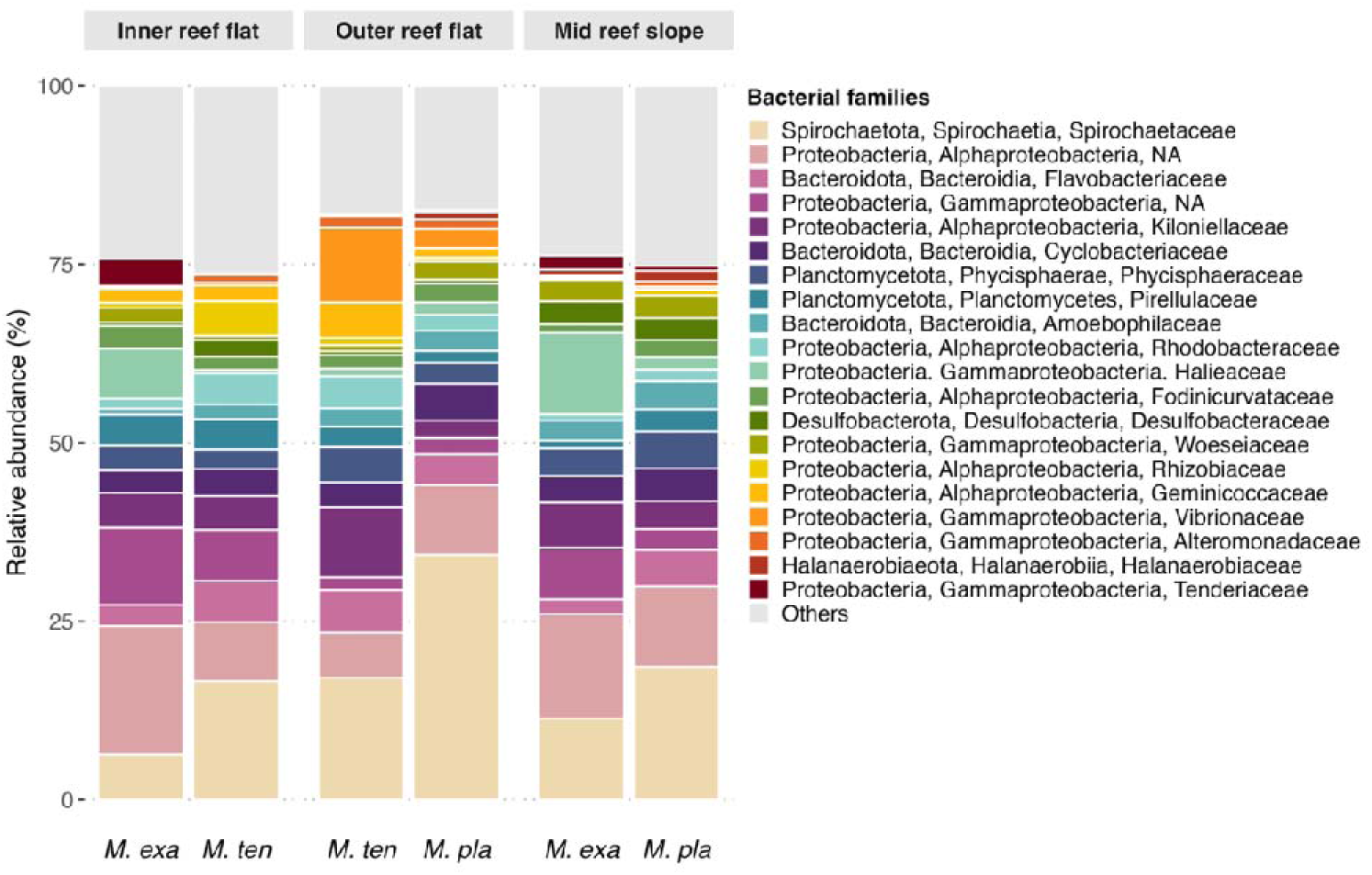
Composition of Prokaryotic assemblages in *Millepora* spp. across host species and reef habitats. The 20 most abundant bacterial families are displayed individually by host species and habitats, while all remaining taxa are grouped in the “Other”. category.

### Symbiont community composition across reef habitats

Despite strong host specificity, Symbiodiniaceae assemblages also exhibited habitat-driven variation within species. At the DIV-level, Symbiodiniaceae assemblages were discriminated by habitat for all three species (sPLS-DA, p < 0.001; Fig. 5A-C). In *M. exaesa*, habitat differences were driven by an enrichment of *Symbiodinium* A16d and A16i in the mid reef slope and X4808043_A in the inner reef flat. Similarly, *M. tenera* in the inner reef flat was defined by a set of host-specific DIVs (i.e., A16bl, A16bm, A16bn and A3bh), aligning with habitat-specific ITS2 profiles including A16bo.A16bl.A16bp in G3 and G5 and A16bl.A16bm.A16.A16bn.A3.A3a in G1, G2, G4 and G6. For *M. platyphylla*, the mid reef slope was discriminated from the outer reef flat by A16h, A3bh and A16bg DIVs, which contributed to the dominant A16. A16bg. A16d. A3. A16h. A3a profile in G16 and G17 (Figs 2 & 5C).

**Figure 5.**
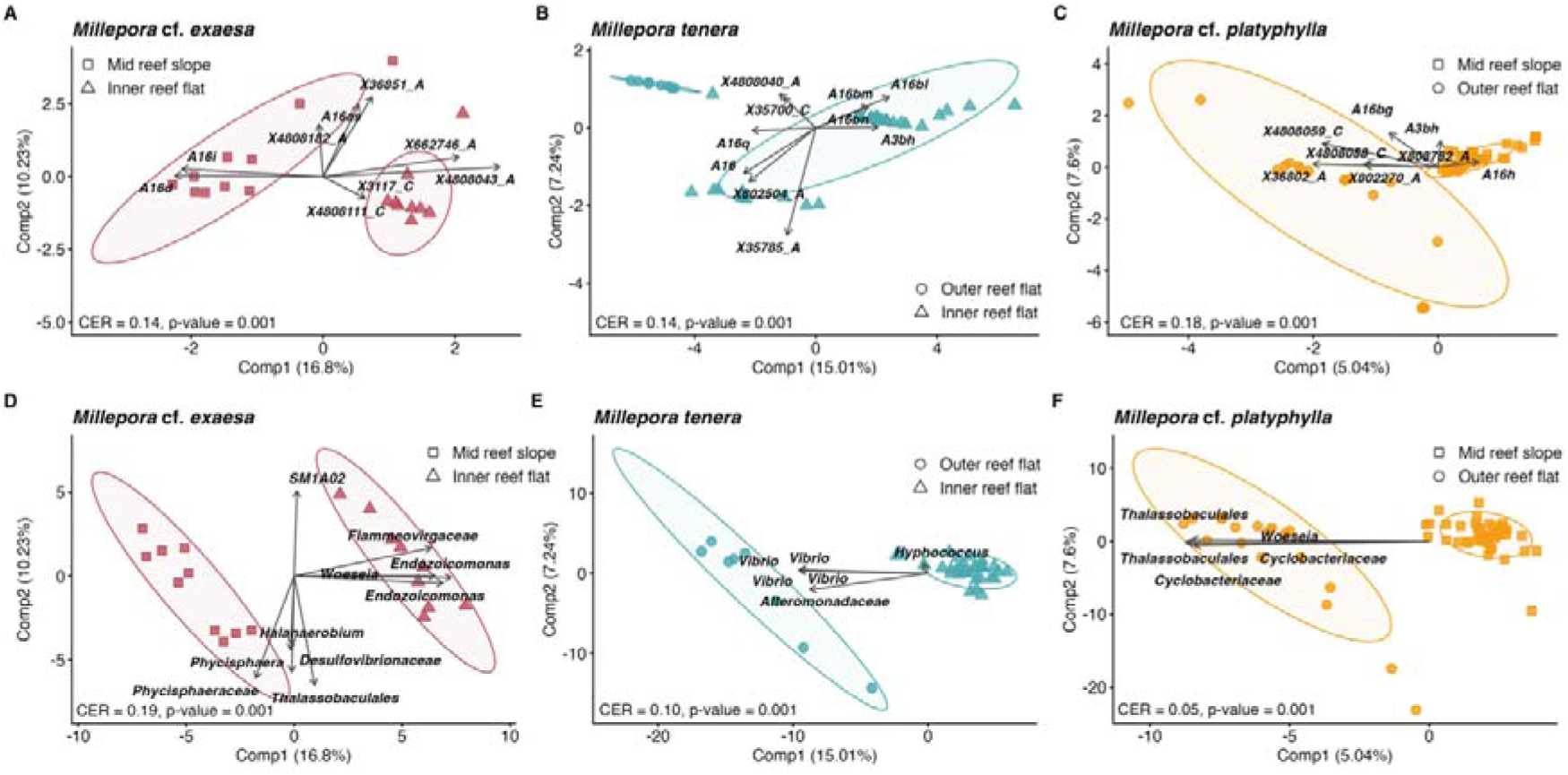
Habitat discrimination based on Symbiodiniaceae and prokaryotic assemblages. sPLS-DA biplots of **(A-C)** ITS2 sequences (DIVs) and **(D-E)** ASVs illustrating the discrimination of habitat for each fire coral species. Points represent individual samples, while arrows indicate the top 5 contributing variables for each species discrimination. Count data were CLR-transformed and scaled. Classification error rate (CER) and pvalue were determined by cross-model validation and permutation tests (nperm = 999). Ellipses represent the spread of the data points at a 95% confidence level.

Within the core microbiome, half of the ASVs (50.4%) were unique to a specific host–habitat combination, while only 3% were shared across all hosts and habitats, highlighting niche specialization. Accordingly, habitat significantly discriminated prokaryotic assemblages (sPLS-DA, p < 0.001; Fig. 5D-F). In *M. exaesa,* habitat discrimination was driven by *Endozoicomonas* and *Woesia* in the inner reef flat, and Phycisphaeraceae in the mid reef slope. In *M. tenera*, several *Vibrio* spp. discriminated the outer reef flat, while *Hyphococcus* characterized the inner reef flat. In *M. platyphylla*, ASVs that discriminated the outer reef flat included unclassified Thalassobaculales and Cyclobacteriaceae. Notably, the Vibrionaceae were enriched in the outer reef flat, both in *M. tenera* (10.5%) and *M. platyphylla* (2.6%; Fig. 4), and core ASVs belonging to *Vibrio*, *Photobacterium*, and *Catenococcus* (Vibrionaceae) were exclusive to *M. tenera* from the outer reef flat (Fig. S7).

### Within-genotype symbiont co-dispersion and co-variation

We then explored to what extent host genotypes influenced co-dispersion and co-variation between symbiont compartments. A positive global correlation was observed between Symbiodiniaceae and prokaryotic within-genotype dispersions (Pearson correlation test, r = 0.48, p = 0.03; Fig. 6A). However, this correlation was driven by host species rather than a true co-dispersion between symbiotic compartments. This pattern likely reflects a Simpson’s paradox artifact, supported by linear models in which the dispersion of one symbiotic group was influenced by species alone and not by the dispersion of the other symbiotic group (LM, Species: p < 0.05; Table S11). Species-level regressions further illustrate this pattern (Fig. 6A). Overall, *M. platyphylla* genotypes showed higher microbiome dispersion than *M. tenera*, despite similar spatial distances between clones (Fig. S1; Fig. 6B; Wilcoxon test, p = 0.23), suggesting that microbiome dispersion constitute a biological trait of the host species.

**Figure. 6.**
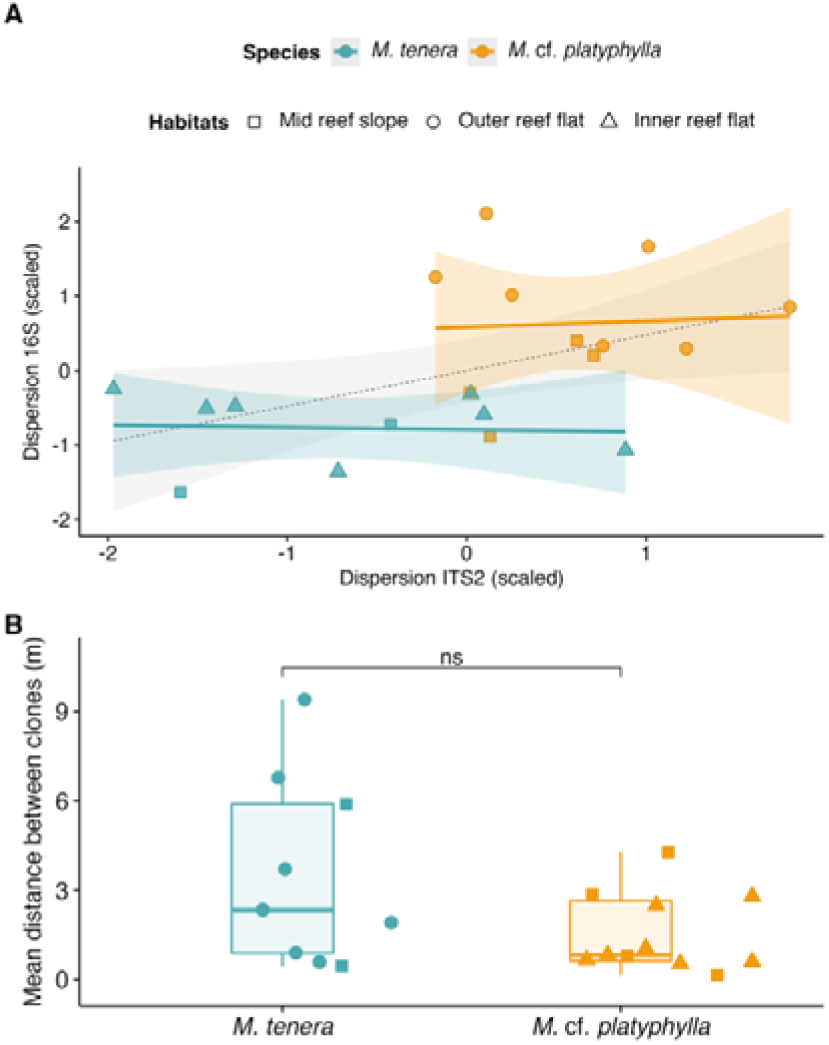
Within-genotype microbiome co-dispersion and spatial distance of *Millepora* species. **(A)** Pearson’s correlation between Symbiodiniaceae and prokaryotic dispersions at the host genotype level. Within-genotype dispersions were calculated from Euclidean matrices of CLR-transformed abundances and scaled within each species to account for species-specific multivariate scales. Statistical testing for the relation between Symbiodiniaceae and prokaryotic dispersions and host species can be found in Table S11 **(B)** Mean spatial distances between clones within each genotype to assess the influence of spatial proximity on microbiome co-dispersion. Statistical differences were evaluated using a non-parametric Wilcoxon test due to the non-normal distribution of the data.

Multi-omic integration (DIABLO) further identified correlated key symbionts that discriminated host genotypes (Figs. 7 & 8). While some genotypes showed defined and congruent symbiotic signature across clones, others tended to have more overlapping microbiomes. For instance, *M. tenera* genotypes G2 and G5 from the inner reef flat showed low multidimensional distances, suggesting a high degree of co-variation between symbiotic compartments (Fig. 7A). G5 microbiome was characterized by *Symbiodinium* A16bp and A16bo with unclassified Kiloniellaceae and Cyclobacteriaceae. G2 harbored *Symbiodinium* A16bl, A16bn and 16bm and *Rubribacterium,* which were shared with G1 and G4, hindering their clear discrimination (Figs. 7A & 8A).

**Figure 7.**
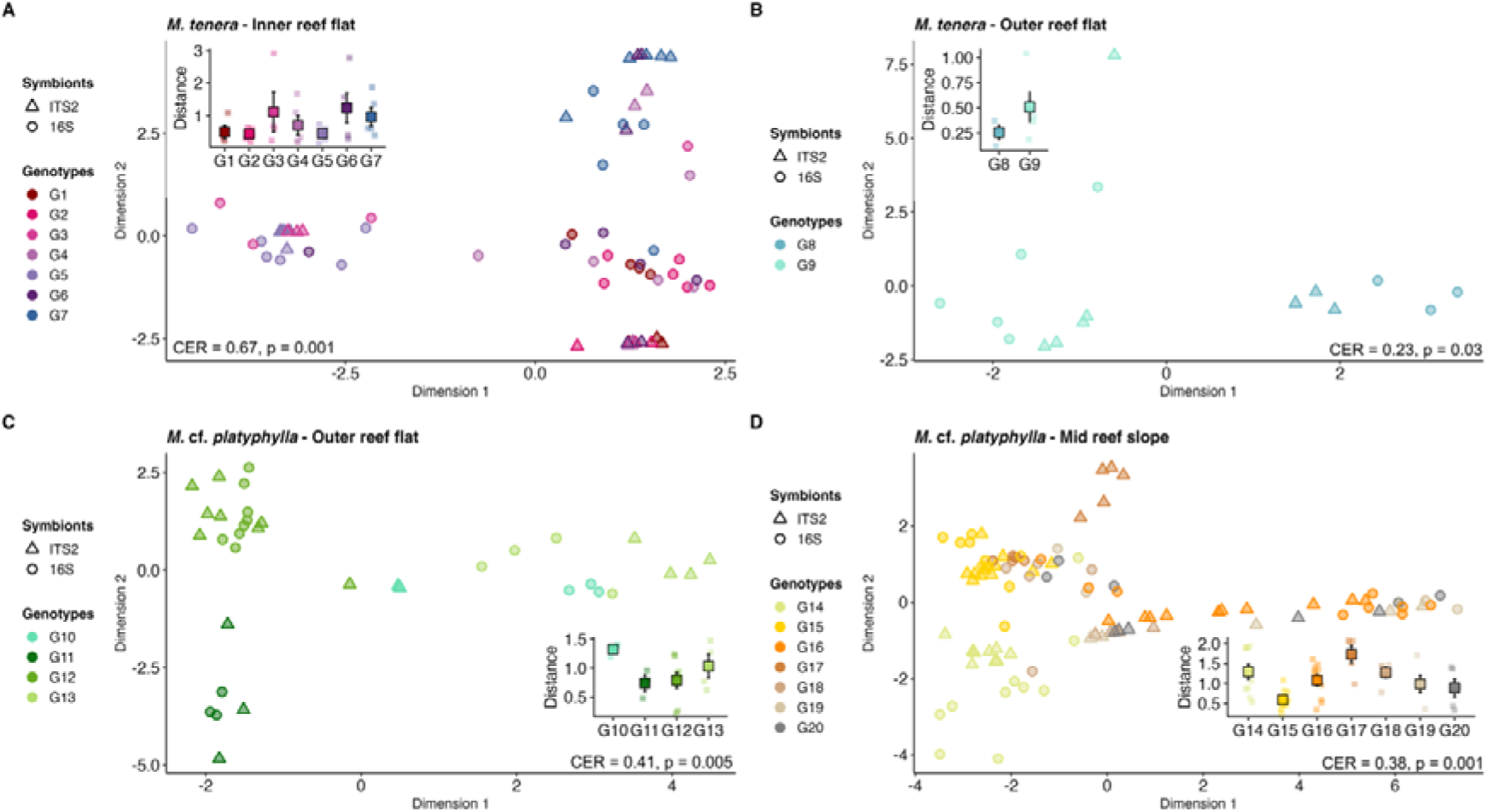
Host genotype discrimination based on the integrated Symbiodiniaceae and Prokaryotic assemblages. Score plots from DIABLO (multiblock sPLS-DA) performed on Symbiodiniaceae ITS2 sequences and prokaryotic ASVs between host genotypes of **(A)** *M. tenera* in the inner reef flat, **(B)** *M. tenera* in the mid reef slope, **(C)** *M.* cf. *platyphylla* in the outer reef flat and **(D)** *M.* cf. *platyphylla* in the mid reef slope. Samples from each dataset are overlaid with a symbol for each symbiotic assemblage and colored by host genotype. The classification error rate (CER) represents the proportion of misclassification. CER and pvalues were determined by cross-model validation and permutation tests (nperm = 999). Insets show mean (± se) multidimensional distance per genotype in the integrative DIABLO space where individual points reflect colony-level concordance between Symbiodiniaceae and prokaryotic communities, calculated as Euclidean distances between paired ITS2 and 16S latent scores.

**Figure 8.**
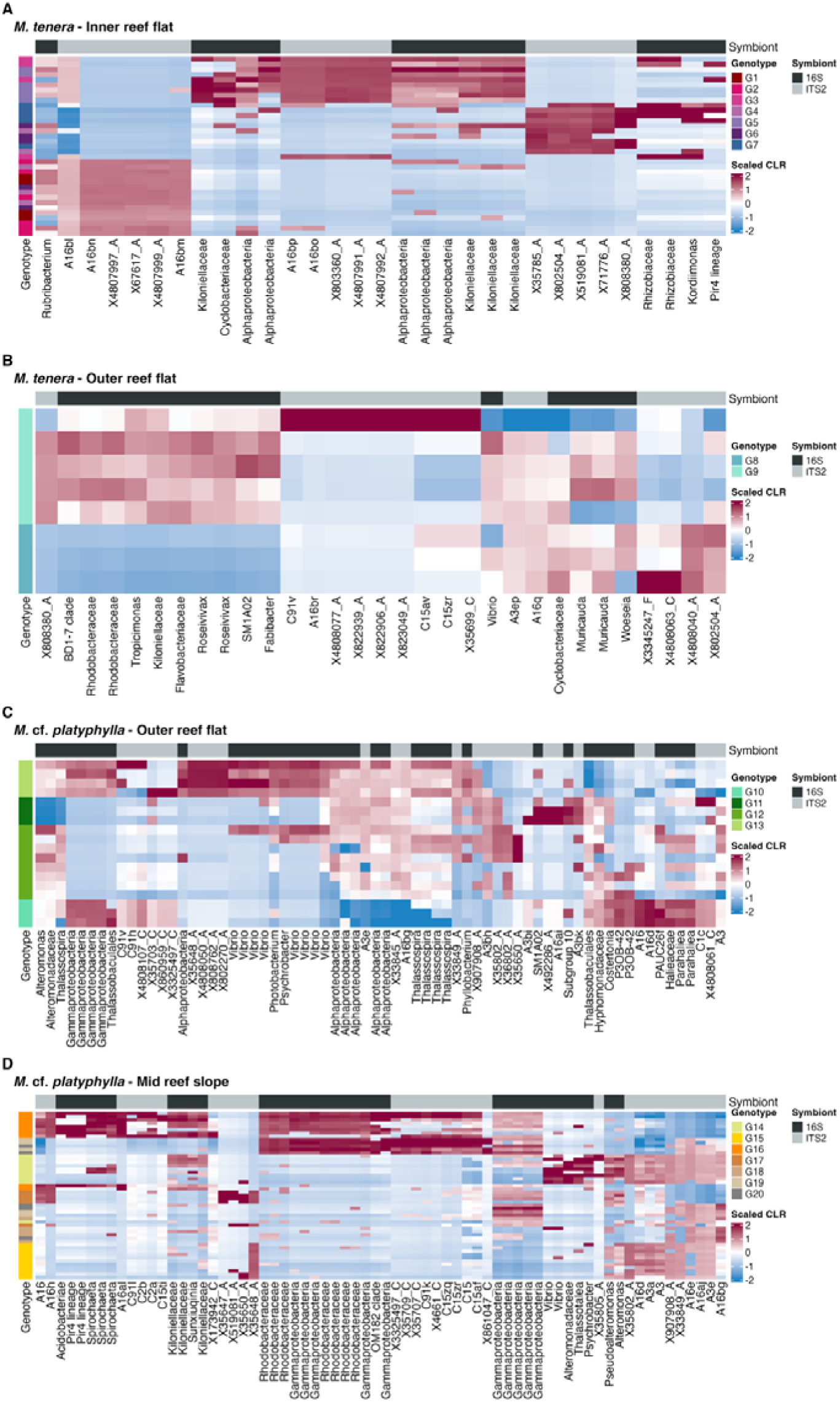
Discriminant and co-varying symbionts contributing to host genotype classification. DIABLO heatmaps illustrating the co-variation patterns between specific Symbiodiniaceae and prokaryotic symbionts that drive the discrimination of host genotypes for **(A)** *M. tenera* in the inner reef flat, **(B)** *M. tenera* in the mid reef slope, **(C)** *M*. cf. *platyphylla* in the outer back reef, and **(D)** *M.* cf. *platyphylla* in the mid reef slope. Heatmaps display the top 10 contributing variables (Symbiodiniaceae DIVs and bacterial ASVs) based on their absolute maximum importance (value of their coefficients) across all components. Selected symbionts (columns) and host genotypes (rows) are clustered based on their CLR-transformed abundances using hierarchical clustering (Euclidean distances).

In the outer reef flat, *M. tenera* genotypes G8 and G9 displayed distinct microbiome signatures (Fig. 7B), with G8 discriminated by *Symbiodinium* X4808040_A X802504_A, *Muricauda* and *Woesia* compared to G9 by *Symbiodinium* X808380_A, *Roseivivax* and *Tropicimonas* (Fig. 8B). Similarly, *M. platyphylla* genotypes in the outer reef flat clearly clustered apart, with a greater congruence between the two symbiotic compartments for G11 and G12 (Fig. 7C). G11 was discriminated by *Symbiodinium* A3bk and A16ai co-varying with Subgroup 10 compared to G12 by *Symbiodinium* A3, *Alteromonas* and *Thalassospira* (Fig. 8C). In the mid reef slope, *M*. *platyphylla* G15 exhibited the strongest microbiome co-variation (Fig. 7D), and was characterized by *Symbiodinium* X_35648A, A16d, A3 and A3a co-varying with *Pseudoalteromonas* and *Alteromonas* (Fig. 8D).

## DISCUSSION

Our study reveals that host genetic identity was the strongest determinant of both Symbiodiniaceae and prokaryotic community structure, while reef habitat played a significant but secondary role. Our integrative approach revealed strong genotypic variability, with some genotypes showing well defined and highly conserved microbiome signatures, and others showing greater microbiome plasticity and overlapping symbiotic communities. For some genotypes, both symbiotic compartments co-varied across clones, suggesting that host genotype can jointly constrain Symbiodiniaceae and prokaryotic assemblages. These findings emphasize that host-symbiont dynamics are not uniform at the inter- and intra-specific levels, highlighting the importance of considering fine-scale host genetic diversity to understand coral holobiont structure and resilience capacity.

Host genetic identity at both the species and subspecies levels was the primary driver of Symbiodiniaceae and prokaryotic assemblages. The consistency of specific ITS2 type profiles across coral clones demonstrates strong specificity and fidelity between the host and their photosymbionts, corroborating numerous studies [21, 23, 27, 28]. The weak overlap in core and discriminating ASVs between species further supports some degree of host specificity [6, 21, 51], likely governed by species-specific skeletal morphology, micro-environmental niches, and molecular recognition [51, 52]. Host-symbiont fidelity might result from vertical symbiont acquisition mode [2, 15, 27]. *Millepora* spp. are broadcast spawners releasing medusoids with sexual gametes, yet they can also transmit algal symbionts to their oocytes [53, 54]. This mixed-transmission mode likely ensures the maintenance of beneficial associations through vertical transmitted symbionts while allowing for environmental acquisition at the larval and medusoid stages, potentially conferring an adaptive advantage in changing environments [14, 55, 56].

Host genotypes showed varying degrees of control across symbiotic compartments, with a stronger effect on Symbiodiniaceae than on prokaryotic community structure. However, distinct host genotypes harbored similar algal symbionts (e.g., *M. tenera* G3 and G5), suggesting that photosymbiont plasticity can match variation in host genetic background, thereby broadening the spectrum of holobiont phenotypes [57, 58]. Although little correlation is often reported between symbiotic compartments [25, 26], our integrative approach revealed genotype-specific co-variation between algal and bacterial assemblages. The extent of this co-variation varied among host genotypes, with some showing highly coordinated microbiome composition (e.g., *M. tenera* G2 & G5; Figs 7 & 8), while others maintained more flexible symbiotic associations. Corals dominated by different Symbiodiniaceae genera can show distinct bacterial community patterns [3, 32]. Yet, fire coral genotypes sharing the same ITS2 profiles showed variable degrees of co-variation, suggesting host genotype, rather than Symbiodiniaceae identity, act as the primary constraint on both symbiotic compartments. These patterns suggest that host genotype may coordinate the recruitment of multiple symbiotic partners, creating an integrated symbiotic network upon which selection can act. However, several genotypes showed decoupled Symbiodiniaceae and prokaryotic assemblages (Figs 7 & 8), suggesting that assembly processes partially differ between symbiotic compartments at the genotype level. Whether taxonomic decoupling between symbiotic compartments translates into functional divergence remains to be determined, and future work incorporating metagenomics or metatranscriptomics will be essential to address this question.

The strength of host genetic control appeared to be species dependent. *M. platyphylla* genotypes exhibited higher diversity of Symbiodiniaceae sequences and higher relative abundance of background symbionts, including *Cladocopium*. In addition, *M. platyphylla* genotype showed greater microbial dispersion regardless of spatial distances between clones, indicating more plastic microbial assemblages within genotypes (Fig. 6). Partnerships with multiple Symbiodiniaceae genera may increase the potential for symbiont shuffling under environmental changes [23], as the capacity of background symbionts to become dominant often depends on their initial abundance, prior disturbances [87]. Given that microbiome regulation differs between host species [29], our results suggest that *M. tenera* imposes stronger genetic filtering over symbiont assembly, while *M. platyphylla* exhibits greater plasticity in both Symbiodiniaceae and prokaryotic assemblages.

The Symbiodiniaceae assemblages of fire corals were dominated by *Symbiodinium* spp. [4], aligning with observations from the South Pacific and Western Indian oceans [27, 28, 59], while Atlantic fire corals predominantly associate with *Breviolum* spp. [60, 61]. All three sympatric *Millepora* species harbored radiations of *Symbiodinium* A16 and A3 variants, alongside *Cladocopium* C15 and C91 (Fig. 2). In the South Pacific (French Polynesia), *M. platyphylla* hosts A7 radiations [27], which were absent from our dataset, while fire corals from the Western Indian Ocean associate with regionally endemic A16 variants [19, 59], reflecting broader biogeographic differences in locally available symbiont pools [62, 63]. Many Symbiodiniaceae lineages remain undescribed at the species levels, although recent efforts are clarifying taxonomy [64]. A3 and A3a variants likely correspond to the generalist photosymbiont *Symbiodinium tridacnidorum* [64], hosted by fire corals, scleractinian corals (*Acropora cervicornis* [65]) and giant clams [66]. Yet, generalist symbionts can mask cryptic host-specialist lineages, as recently shown for *Cladocopium* C15 in *Porites* spp. [67]. Indeed, while A16 and A3 variants are widely shared across all three *Millepora* species, DIV-level analysis revealed strong variation across species, with *M. exaesa* associating with distinct lineages (A16a, A16j and A16be; Figs 2 & 3) with no overlap with *M. platyphylla* or *M. tenera*. This divergence aligned with host phylogeny (Fig. S1). Whether these host-specific variants represent cryptic *Symbiodinium* species or subspecies that have co-diverged with their hosts remains unresolved and warrants further genetic investigation.

All three fire coral species consistently associated with Spirochaetaceae, Kiloniellaceae and Desulfobacteraceae, across reef habitats (Fig. 4; Fig S6), suggesting evolutionarily maintained functional benefits [16, 18]. Spirochaetaceae (e.g., *Spirochaeta*) dominant in both soft [68] and fire corals [1], are known for anaerobic fermentation of carbohydrates and nitrogen fixation [69, 70]. In the low-oxygen microenvironment of coral skeletons [51, 71], they could produce fermentation byproducts (e.g., hydrogen, acetate), forming syntrophic loops with sulfate-reducing bacteria like *Desulfospira* [71] within the holobiont. High abundances of Spirochaetaceae were recently identified as a signature of stress-resistant genotypes in *Acropora cervicornis* [72], suggesting they may underpin bleaching resilience in *Millepora* [53]. Additional core bacteria may be potentially involved in nitrogen fixation (*Tistlia* [73]), polysaccharide degradation (*Fulvivirga* [74]), DMSP metabolism and antioxydant production (*Muricauda* [75]). While the observation of specific core taxa across species-habitat combinations suggests niche specialization, most bacterial functions are widespread across taxa [1, 9, 12, 76]. Thus, taxonomic divergence in the prokaryotic core assemblages may reflect competitive interactions and/or differences in symbiotic compatibility rather than functional specialization alone. Moreover, some functions provided by bacterial symbionts may overlap with those of algal symbionts (e.g., *Cladocopium* C15), creating a degree of functional redundancy within the holobiont. Such metabolic redundancy across symbiotic compartments may be particularly beneficial during stress events, including bleaching, when the loss of Symbiodiniaceae increases the reliance of the holobiont on bacterial partners to sustain essential metabolic functions [8] .

Symbiodiniaceae and prokaryotic assemblages shifted within each habitat in host-specific ways (Figs 2 & 3), demonstrating that environmental filtering shapes symbiotic communities, although within boundaries imposed by host genetic identity. Inshore reefs often show extreme and variable light and temperature regimes compared to fore reefs [1, 25]. In Reunion Island, freshwater run offs and groundwaters infiltration can further influence water salinity and nutrient levels [77]. However, the contribution of reef habitats remained secondary to host genetics, contrasting with patterns observed in Mo’orea, where *M. platyphylla*, the sole fire coral species present, harbor prokaryotic communities that are primarily structured by environmental variation [1]. Consistent with the negative correlation between species richness and niche breadth [78], the sympatric distribution of three *Millepora* species in Reunion Island may constrain each host to a narrower and more specialized microbial niche. Consequently, each *Millepora* species likely recruits symbionts from locally available pools shaped by habitat conditions, while host-symbiont genetic compatibility acts as the primary filter [79]. However, our observations are based on a single reef site with a complex disturbance history (e.g., marine heat waves, mudslide) which may not reflect species-wide patterns around Reunion Island. In our study, *M. tenera* was encountered only in the lagoon, preventing us from confidently distinguish species-specific from habitat-driven patterns.

The establishment of symbiotic partnerships within defined habitats likely provide functional advantages tailored to local environmental conditions [1, 25, 80]. In scleractinian corals, *Cladocopium goreaui* (C1) from nearshore reefs exhibits higher photochemical efficiencies during thermal stress compared to forereef populations, supporting greater holobiont thermotolerance [81]. Our result suggests similar adaptative or acclimatory potential in fire corals. In the inner reef flat, *M. tenera* hosted habitat-specific Symbiodiniaceae assemblages across multiple host genotypes (Fig. 2) and was enriched in *Rubribacterium* and *Inquilinus* (Fig. 8; Fig. S6), taxa characteristic of high-temperature and high-irradiance reefs [80, 82], suggesting a combined contribution of algal and bacterial symbionts to holobiont thermal tolerance. Additionally, in the outer reef flat, we observed a convergence of prokaryotic assemblages between distinct genotypes (i.e., *M. tenera* G8 & G9), suggesting that environmental filtering can override host genetics in challenging habitats [1].

This study demonstrates that host genetic identity is a major determinant of holobiont assembly in sympatric species, while local environmental conditions further modulate symbiotic community composition. Symbiont assemblages, Symbiodiniaceae particularly, were constrained by host genetic boundaries, promoting niche partitioning within reef habitats. Our integrative approach further revealed that some genotypes maintained coordinated algal and bacterial assemblages across clones, suggesting that multiple symbiotic compartments can be structured by common host-genetic filters. Future studies integrating -omic approaches will be essential to determine whether these taxonomic associations translate into functional differences among holobionts [58, 83, 84]. By revealing coordinated genetic control of both algal and bacterial assemblages, our findings show that intraspecific host variation plays a central role in shaping microbiome assembly and holobiont structure.

## DATA AVAILABILITY

Metabarcoding data have been deposited to NCBI Sequence Archive Read under the BioProject accession number PRJNA1527031 with separate SRA accessions for the ITS2 and 16S rRNA datasets. Host mt16S data are available on GenBank under the accession numbers PZ859643-PZ859683. Metadata, microsatellites data and raw ITS2 and 16S rRNA count data can be downloaded from the Zenodo repository <u>10.5281/zenodo.23027540</u>.

Reviewers can access the repository with the following link: https://zenodo.org/records/23027541?token=eyJhbGciOiJIUzUxMiIsImlhdCI6MTc5MjI5NzI3NywiZXhwIjoxNzk4Njc1MTk5fQ.eyJpZCI6IjVmZGIwN2RmLWM0N2EtNGNhZC1iYjg3LWU2ZTMxNWI4ZmY3NiIsImRhdGEiOnt9LCJyYW5kb20iOiIxZDY1NmViMzMyMmMxZjFkNDRjNzgxNzdjZGUwMzAxNSJ9.y4aeydnUWmiXya7e4pvRK_2zdleU5EQ19JntWz3ip7XJ9imzmkUNQJeN16h2z1QvofAm_qlpzJZc-rRXbzCJzQ

## CODE AVAILABILITY

R scripts used for the analysis are available at the Zenodo repository 10.5281/zenodo.23027540.

## Acknowledgments

The authors would like to thank Edouard Jobet for facilitating lab work, Sophie Bureau for field work and Lucas Venegas for 16S rRNA metabarcoding data processing. This work was funded by the Laboratoire d’Excellence “CORAIL” project COMIC project led by CA-FB, and a L’Oréal-Canada grant awarded to CED. CPS was supported by a FRQNT postdoctoral scholarship (Grant number 352800) and CD was supported by a NSERC and FRQNT postdoctoral scholarships.

## Author contributions

CED, EB and CA-FB. conceived and designed research. CED, AM and CA-FB collected field data. CPS and CED analyzed and interpreted data. CPS, GM, P-LM and CED generated molecular data. CPS wrote the manuscript. All authors revised drafts of the manuscript and gave final approval for publication.

