## Supplementary information for "Host genetics structure Symbiodiniaceae and prokaryotic assemblages and their co-variation in sympatric fire coral species"

**SUPPLEMENTARY METHODS**

***Millepora* species and genotype delineation**

We sequenced the mitochondrial 16S gene to verify the identity of each sampled *Millepora* species, on all *M.* cf*. exaesa* samples and one representative sample per clonal genotype for *M. tenera* and *M.* cf*. platyphylla*. PCR reactions of 25 *µ*L were prepared for each sample containing 2 *µ*L of template DNA, 0.6 *μ*L of each primer [10 *μ*M], 2.5 *μ*L of dNTPs [2 mM], 2 *μ*L of MgCl2 [25 mM], 2.5 *μ*L of buffer [10×], 0.1 *μ*L of TAQ polymerase [5 unit/μl] and 14.7 *μ*L of DNA- free water (Taq DNA Polymerase kit, Qiagen, Hilden, Germany). PCR cycling parameters were set with an initial denaturation step of 1 min at 95°C, followed by 40 cycles of 45 s at 94°C, 1 min at 52°C and, 1 min’10 s at 72°C and a final extension phase of 4 min at 72°C.

Sequence chromatograms were manually inspected to resolve ambiguous N bases when supported by peak quality and alignment consensus with other sequences. mt16S sequences were aligned using MAFFT with default settings [1] within Unipro UGENE software (v52.1; [2]). Additional *Millepora* sequences (three for each species) from Réunion Island [3] were included to unsure the correct species assignation. A sequence of *Apatizanclea divergens* (Hydrozoa, Zancleidae family) was used as an outgroup to root the phylogenetic tree. Maximum likelihood (ML) phylogenetic trees were inferred using IQ-TREE 2 [4], with ultrafast bootstrap (1000 replicates; [5]) and SH-aLRT support values. ModelFinder estimated TPM2u+F+G4 as the best-fit substitution model based on Bayesian Information Criterion (BIC) scores [6]. The resulting ML phylogenetic trees were annotated on R (version 4.5.1).

For clonal genotype identification, a total of 9, 15 and 17 polymorphic microsatellite loci were amplified for *M.* cf*. exaesa*, *M. tenera* and *M.* cf. *platyphylla*, respectively, [7, 8] in five PCRs using the Type-it Multiplex Master Mix (QIAGEN). PCR products were visualized on an Applied Biosystems 3730 capillary sequencer with an internal size standard (GeneScan 500 LIZ) by the GenoScreen plateform (Lille, France). Alleles were manually scored and verified based on amplicon size using GeneMapper v4.0 (Applied Biosystems, USA). Multilocus genotypes (MLGs) were identified using GenClone v2.0 [9]. Genets were defined as identical MLGs across all loci, with the probability of identical genotypes arising from fragmentation set to < 0.001 using with GenAlEx v6.5 [10]. MLGs detected in a single colony were considered unique and assumed to result from sexual reproduction.

**PCR conditions ITS2 and 16S rRNA genes amplification**

Symbiodiniaceae ITS2 amplicons were generated using the primers SYM_VAR_5.8S2 (forward: 5’-TCGTCGGCAGCGTCAGATGTGTATAAGAGACAGGAATTGCAGAACTCCGTGAACC-3’) and SYM_VAR_REV (reverse: 5’: GTCTCGTGGGCTCGGAGATGTGTATAAGAGACAGCGGGTTCWCTTGTYTGACTTCATGC-3’) primers (Hume et al 2018). Prokaryotic 16S rRNA gene amplicons were generated using the primers 519F (forward: 5’- ACACTCTTTCCCTACACGACGCTCTTCCGATCTCAGCMGCCGCGGTAA-3’) and 803R (reverse:GTGACTGGAGTTCAGACGTGTGCTCTTCCGATCTCTACCRGGGTATCTAATCC-3’) with Illumina adaptor overhangs underlined.

PCR conditions for ITS2 amplification were 98°C for 2 min, followed by 30 cycles of 98°C for 10 s, 58°C for 30 s and 72°C for 30 s with a final extension at 72°C for 2 min. For 16S rRNA amplification, PCR conditions differ only with 60°C as melting temperature. Successful amplification was verified on a 2% agarose gel. DNA samples were then purified using magnetic beads (AMPure XP, Beckman Coulter) at 70% volume and prepared at a concentration of 5-10 ng/µL. Indexing adaptors (i.e., Illumina dual index adaptors) were added via PCR reactions of 25 µL using the same amplification conditions as above but for 9 cycles. Indexed PCR products were purified using magnetic beads and quantified using a Nanodrop 2000 Spectrophotometer (Thermo Scientific). Samples of amplified DNA were submitted for sequencing to CHUL of Québec, Université Laval, and paired-end sequenced (2x250bp) on the Illumina NovaSeq 6000 platform.

**SUPPLEMENTARY FIGURES**

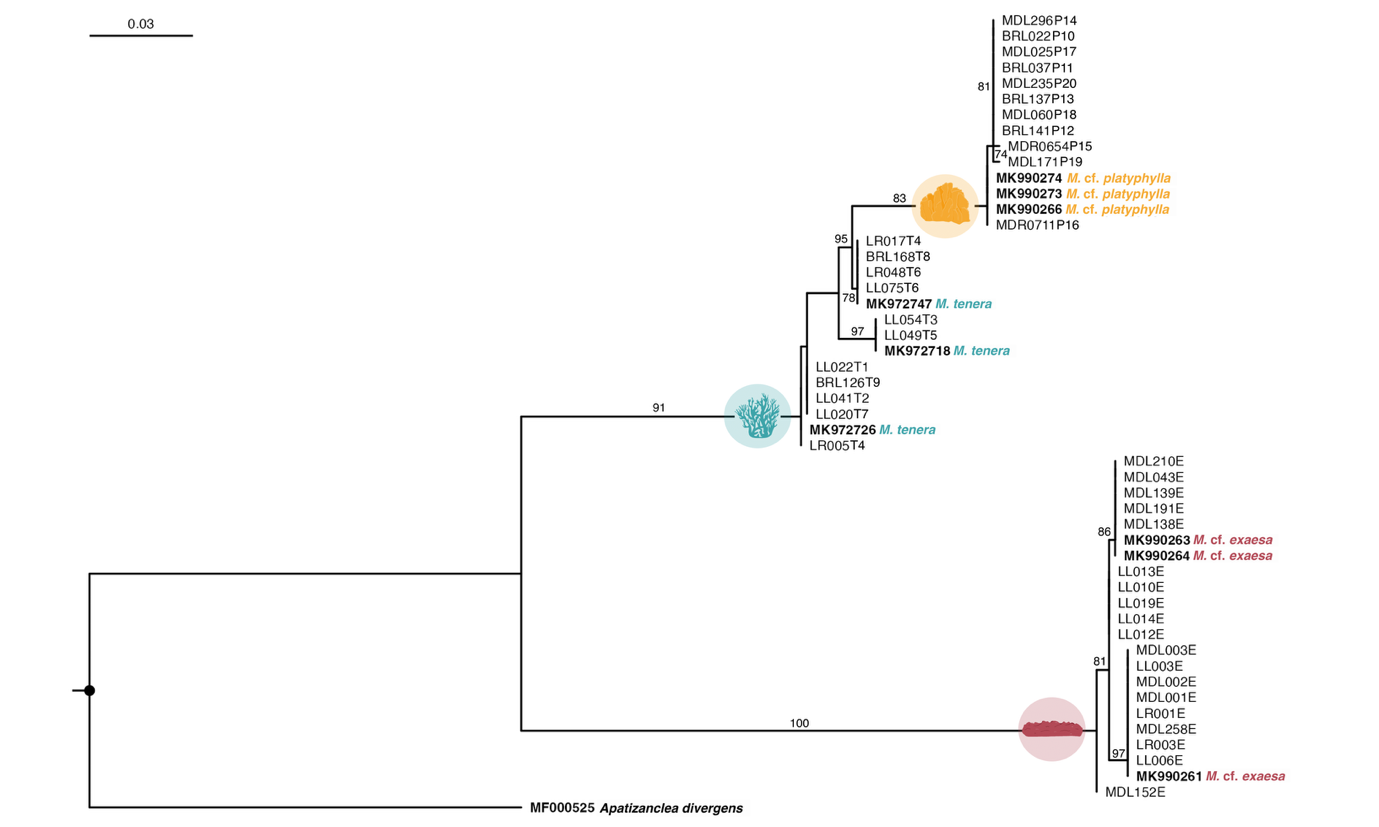

**Fig. S1.** **Millepora species phylogenetic reconstruction**. Maximum Likelihood reconstruction of the mt16S sequencing data from the three Millepora species: Millepora cf. platyphylla, Millepora tenera and Millepora cf. exaesa. In sample names, end letters “P”, “T” and “E” corresponds to the species, M. cf. platyphylla, M. tenera and M. cf. exaesa, respectively, with the last digits corresponding to the host genotype. Three sequences from Boissin et al. 2022 [3] were added for each species as reference to confirm species clusters, alongside the outgroup sequence for Apatizanclea divergens, all available on NCBI GenBank. See Data availability statement. Branch support bootstrap values are indicated when > 70.

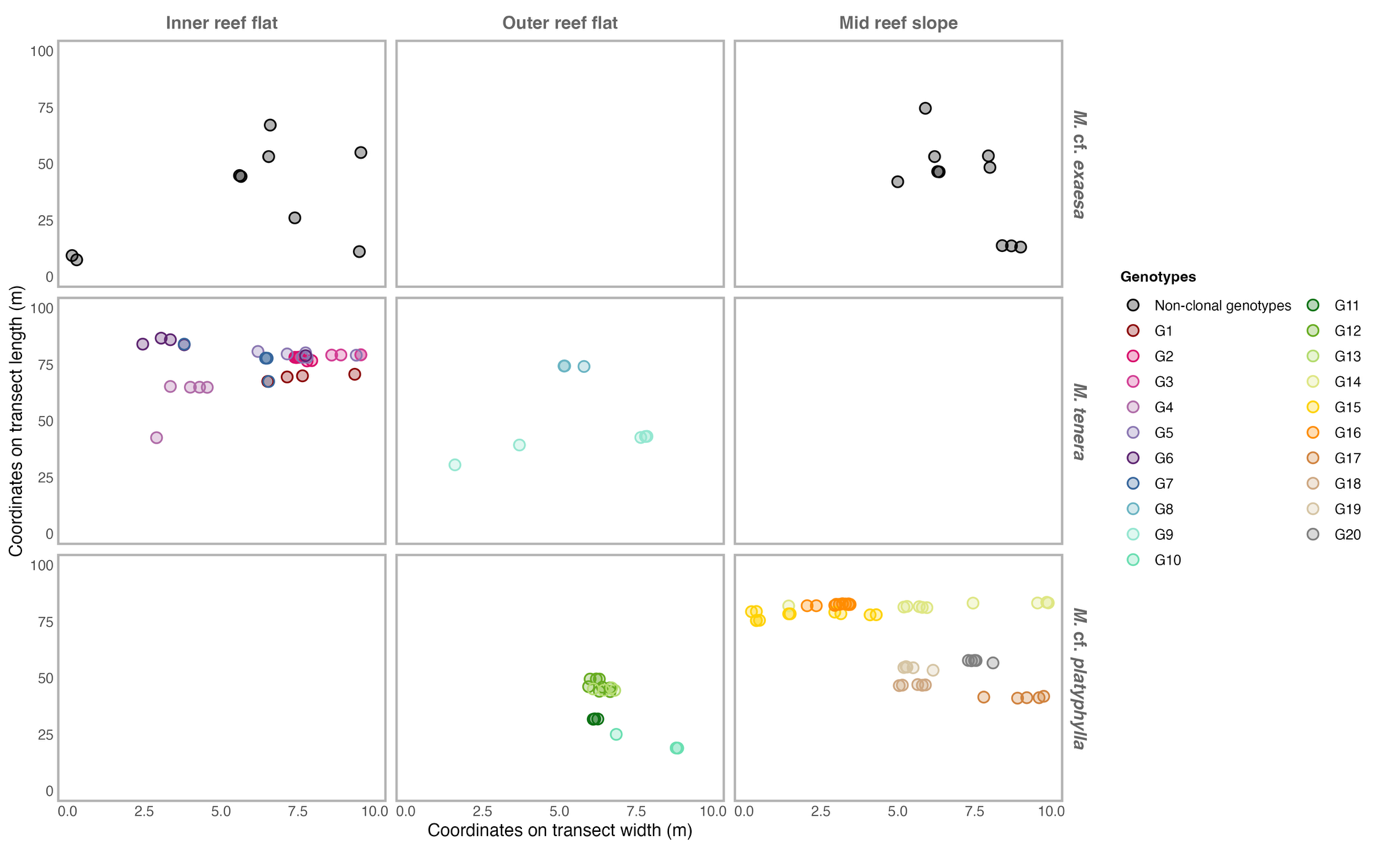
**Fig. S2.** **Spatial distribution of Millepora species at St-Leu, Reunion Island**. Each colony was mapped along belt transect with its x and y coordinates corresponding to its position on the transect width and length. M. cf. exaesa, M. tenera and M. cf. platyphylla occured in three adjacent habitats: the inner reef flat, outer reef flat and mid reef slope. Clonal genotypes were observed for M. tenera and M. cf. platyphylla.

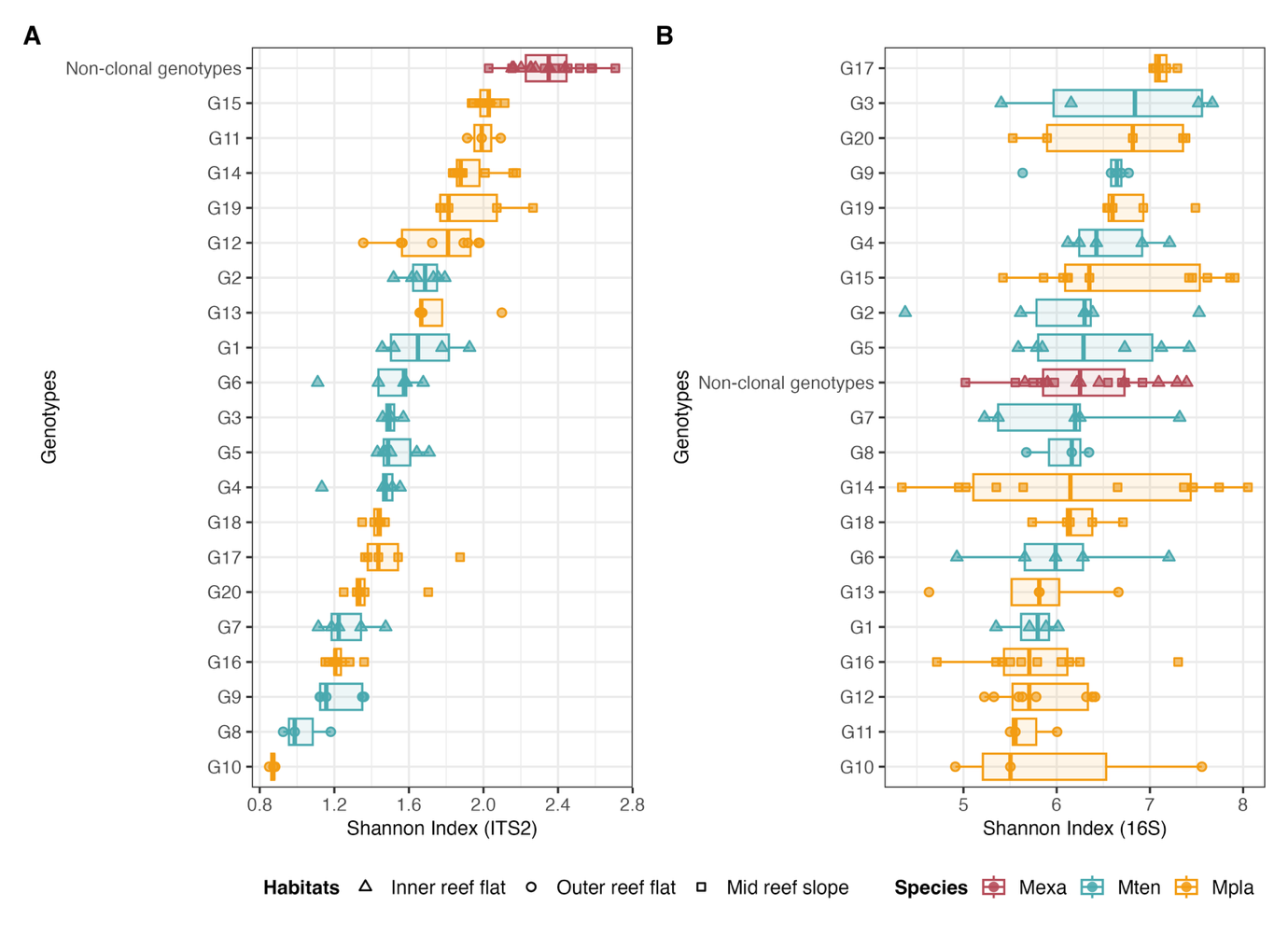

**Fig. S3.** **Microbial alpha-diversity of Millepora species across host genotypes and reef habitats.** Shannon diversity index of individual colonies of **(A)** Symbiodiniaceae DIVs (ITS2) and **(B)** prokatyotic ASVs (16S rRNA). Genotypes within each habitat and species are ranked by decreasing diversity values. Results for statistics can be found in Table S4.

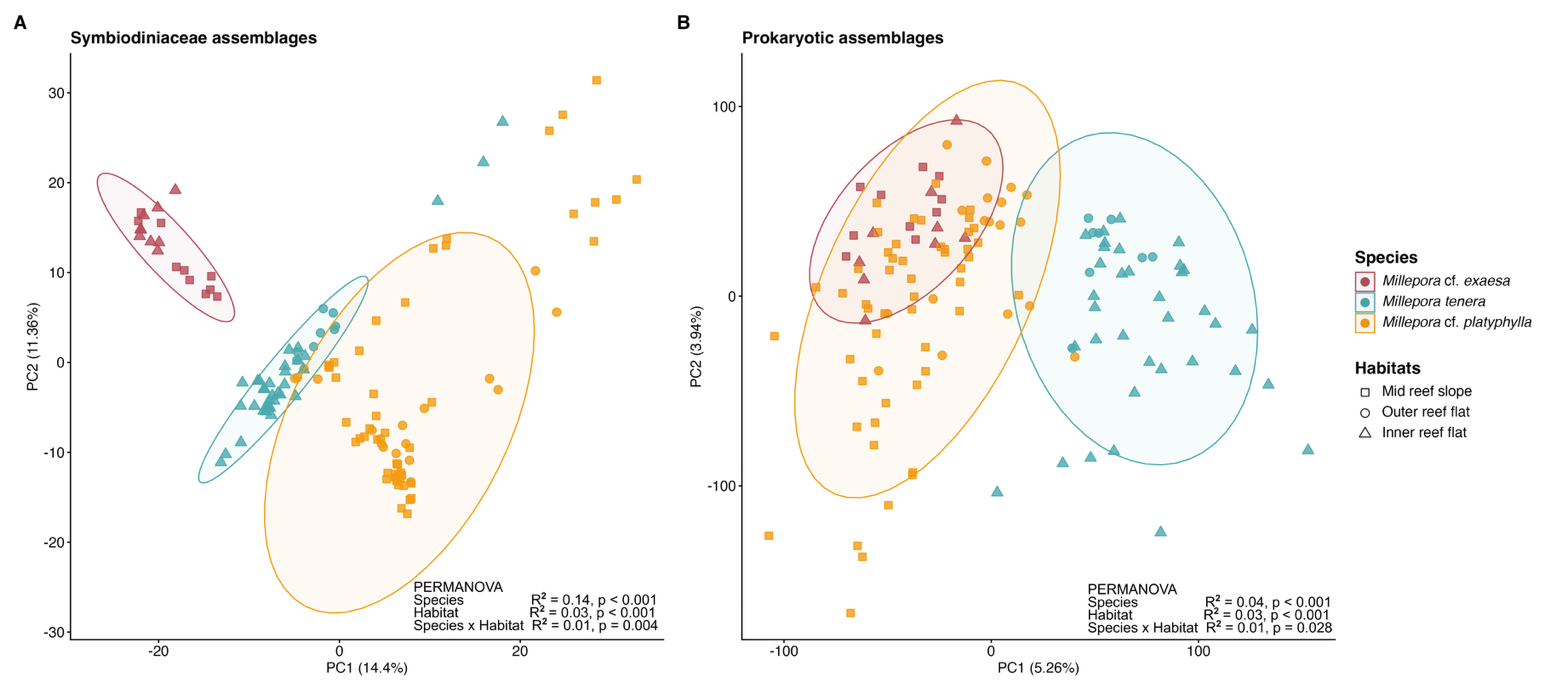
**Fig. S4.** **Microbial beta-diversity of Millepora species across reef habitats.** PCA score plots performed on **(A)** Symbiodiniaceae and **(B)** Prokaryotic assemblages using CLR-transformed abundances. Ellipses represent the spread of the data points at a 95% confidence interval. Compositional differences across species and habitats were tested by PERMANOVA (nperm = 999). Pairwise comparisons can be found in Tables S5 & S6 and betadispersion analysis in Table S7.

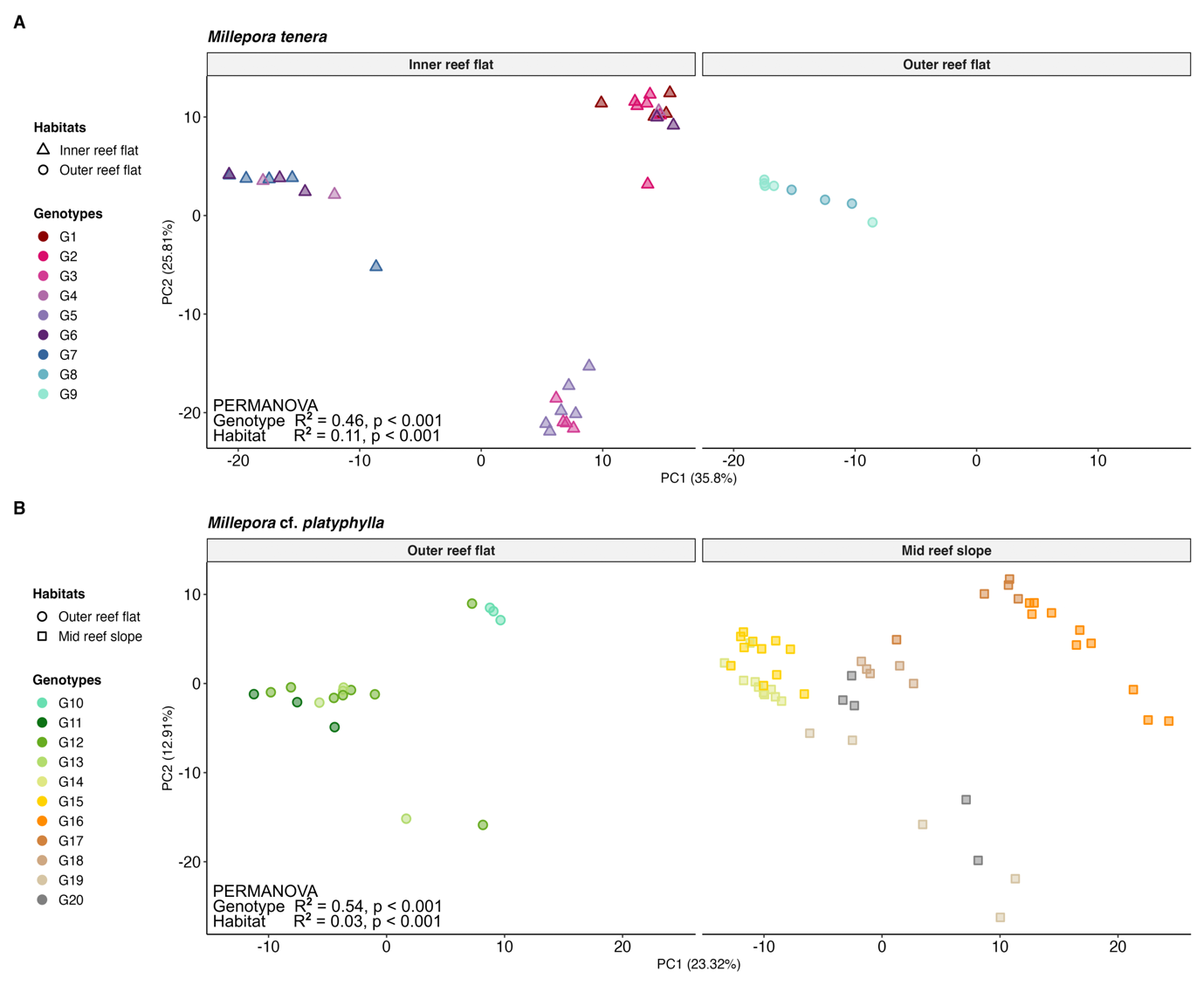

**Fig. S5.** **Symbiodiniaceae beta-diversity of Millepora species across host genotypes and reef habitats.** PCA score plots performed on **(A)** M. tenera and **(B)** M. cf. platyphylla using CLR-transformed abundances. Sample points within each habitat were plotted in distinct facet for better visualization, although a single PCA were performed per host species. Contribution of host genotypes and habitats on Symbiodinaceae assemblages were tested by PERMANOVA (nperm = 999). Pairwise comparisons can be found in Table S8 and betadispersion analysis in Table S10.

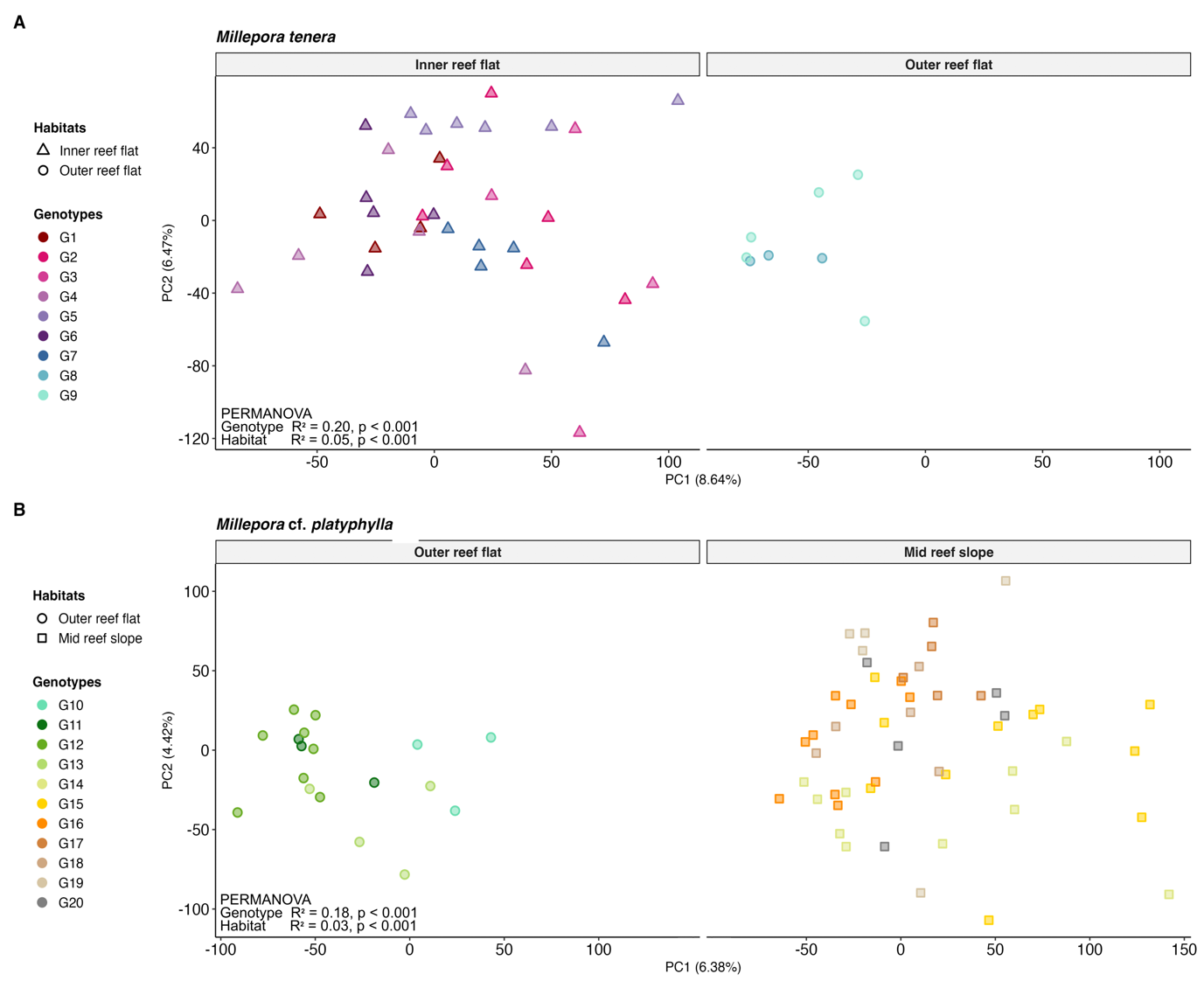

**Fig. S6.** **Prokaryotic beta-diversity of Millepora species across host genotypes and reef habitats.** PCA score plots performed on **(A)** M. tenera and **(B)** M. cf. platyphylla using CLR-transformed abundances. Sample points within each habitat were plotted in distinct facet for better visualization, although a single PCA were performed per host species. Contribution of host genotypes and habitats on Prokaryotic assemblages were tested by PERMANOVA (nperm = 999). Pairwise comparisons can be found in Table S9 and betadispersion analysis in Table S10.

**Fig. S7. Core taxa in the Prokaryotic assemblages of Millepora species across reef habitats.**
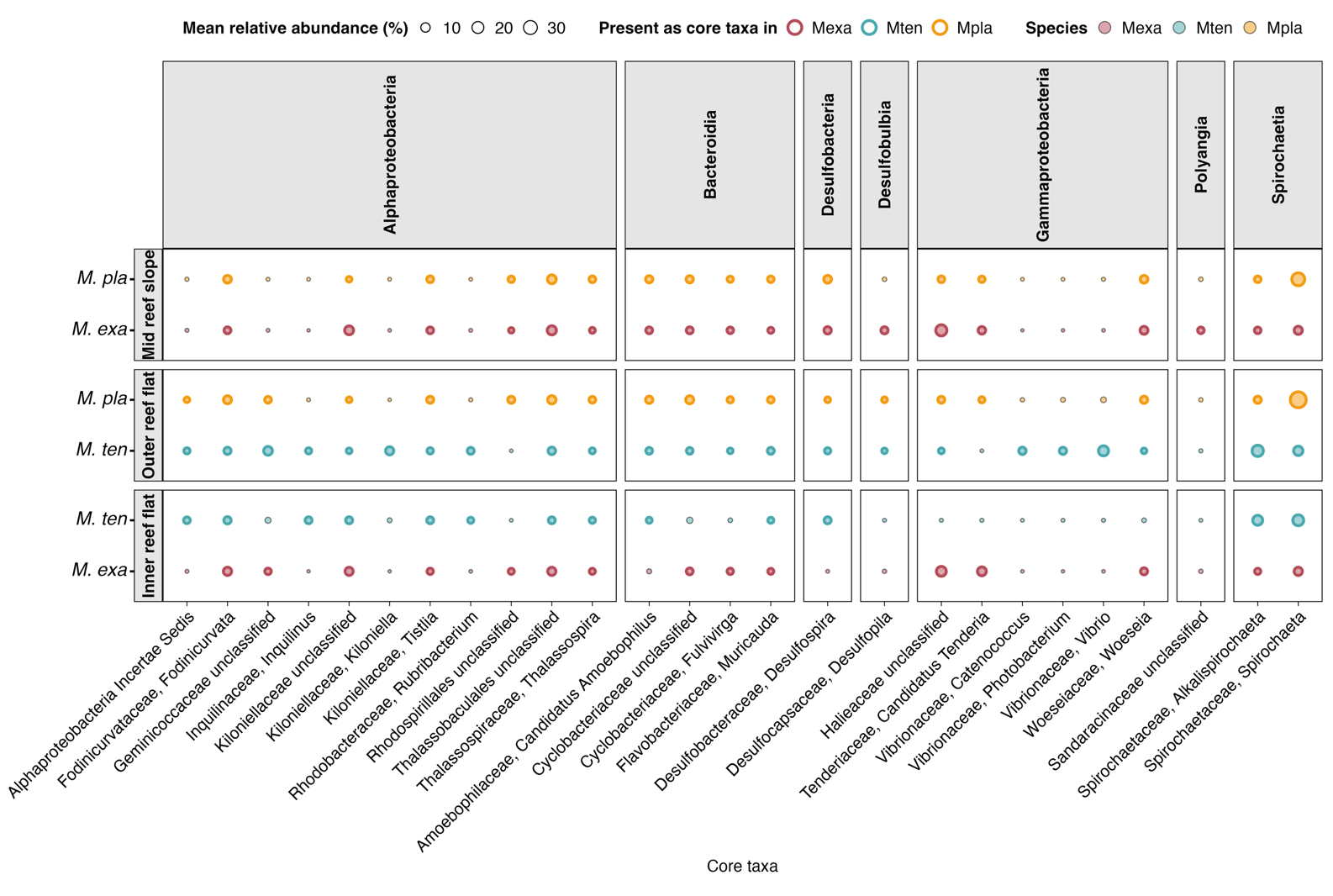
Core members within each host species and habitat combination were identified based on a 80% occurence threshold. Prokaryotic taxa were present in all species and habitat but not necessarily as core members.

**SUPPLEMENTARY TABLES**

Available as .csv files at the Zenodo repository [10.5281/zenodo.23027540](https://doi.org/10.5281/zenodo.23027540).

**Table S1**: Sample metadata with the number of reads from the ITS2 and 16S rRNA datasets, the number of ITS2 defining intragenomic variants (DIVs) and type profiles, the number of amplicon sequence variants (ASVs).

**Table S2:** Raw microsatellite data used for multilocus genotyping

**Table S3:** Raw count data of ITS2 DIVs and type profiles in each sample

**Table S4:** Raw count data of ASVs in each sample

**Table S5:** **Statistical testing for the influence of *Millepora* spp. host identity and habitat on microbial alpha-diversity (Shannon index) of both symbiotic compartiments.** At the species-level for the Symbiodiniaceae assemblages, Kruskal-Wallis results for the effects of species, habitats and their combinaiton followed by post-hoc Dunn’s tests with BH correction. For the prokaryotic assemblages, ANOVA results of linear models with species, habitat and their interaction as fixed factors. At the subspecies level, ANOVA results of linear models testing the effects of habitat and host genotype nested within habitat, separately for *M. tenera* & *M.* cf . *platyphylla*.

| **Symbiodiniaceae assemblages** | *Source of variation* | *df* |  | *Chi^2^* | *pvalue* |
| --- | --- | --- | --- | --- | --- |
|  | Species | 2 |  | 54.70 | **< 0.001** |
|  | Habitats | 2 |  | 7.13 | **0.03** |
|  | Species x Habitats | 5 |  | 60.75 | **< 0.001** |
|  | *Pairwise comparisons*  Mexa : IRF vs. MRS  Mten : IRF > ORF  Mpla : ORF vs. MRS  MRS : Mexa > Mpla  ORF : Mpla > Mten  IRF : Mexa > Mten |  |  |  | 0.07  **< 0.001**  0.93  **< 0.001**  **0.01**  **< 0.001** |
|  |  | *df* | *SS* | *F* | *pvalue* |
| *M. tenera* | Habitat | 1 | 0.83 | 37.90 | **< 0.001** |
|  | Genotype (nested) | 7 | 0.67 | 4.35 | **0.002** |
|  | Residuals | 34 | 0.75 |  |  |
| *M.* cf. *platyphylla* | Habitat | 1 | 0.01 | 0.29 | 0.59 |
|  | Genotype (nested) | 9 | 7.50 | 38.3 | **< 0.001** |
|  | Residuals | 58 | 1.26 |  |  |
| **Prokaryotic assemblages** | *Source of variation* | *df* | *SS* | *F* | *pvalue* |
|  | Species | 2 | 0.03 | 0.02 | 0.97 |
|  | Habitats | 2 | 3.23 | 2.55 | 0.07 |
|  | Species x Habitats | 1 | 3.17 | 4.99 | **0.03** |
|  | Residuals | 125 | 79.32 |  |  |
|  | *Pairwise comparisons*  Mexa : IRF vs. MRS  Mten : IRF vs. ORF  Mpla : ORF < MRS  MRS : Mexa vs. Mpla  ORF : Mpla vs. Mten  IRF : Mexa vs. Mten |  |  |  | 0.21  0.79  **0.004**  0.20  0.14  0.28 |
| *M. tenera* | Habitat | 1 | 0.05 | 0.08 | 0.78 |
|  | Genotype (nested) | 7 | 3.43 | 0.84 | 0.56 |
|  | Residuals | 34 | 19.86 |  |  |
| *M.* cf. *platyphylla* | Habitat | 1 | 5.34 | 7.61 | **0.008** |
|  | Genotype (nested) | 9 | 8.97 | 1.42 | 0.20 |
|  | Residuals | 58 | 40.70 |  |  |

**Table S6. Contribution of host species and habitats on microbial community composition.** Permutational Multivariate Analysis of Variance (PERMANOVA, nperm = 999) testing the effects of host species and reef habitat (Inner Reef Flat, IR; Outer Reef Flat, OR; and Mid Reef Slope, MRS) on CLR-transformed abundances. Analyses were performed using Marginal Sum of Squares (by = "margin") to account for the unbalanced experimental design, ensuring that the order of covariates does not bias the explained variance. Separate models were run to assess the main effects of species and habitat, and their interaction (Species x Habitats).

| **Symbiodiniaceae assemblages** | *Source of variation* | *df* | *R2* | *F* | *pvalue* |
| --- | --- | --- | --- | --- | --- |
|  | Species | 2 | 0.14 | 11.61 | **0.001** |
|  | Habitats | 2 | 0.03 | 2.46 | **0.001** |
|  | Species x Habitats | 2 | 0.01 | 2.48 | **0.004** |
|  | *Pairwise comparisons*  Mexa : IRF vs. MRS  Mpla : ORF vs. MRS  Mten : IRF vs. ORF  MRS : Mexa vs. Mpla  ORF : Mpla vs. Mten  IRF : Mexa vs. Mten |  | 0.14  0.03  0.07  0.14  0.18  0.21 | 2.79  1.96  3.29  4.83  2.56  10.95 | **0.005**  **0.019**  **0.006**  **0.001**  **0.001**  **0.001** |
| **Prokaryotic assemblages** | *Source of variation* | *df* | *R2* | *F* | *pvalue* |
|  | Species | 2 | 0.04 | 2.98 | **0.001** |
|  | Habitats | 2 | 0.03 | 1.90 | **0.001** |
|  | Species x Habitats | 2 | 0.01 | 1.27 | **0.028** |
|  | *Pairwise comparisons*  Mexa : IRF vs. MRS  Mpla : ORF vs. MRS  Mten : IRF vs. ORF  MRS : Mexa vs. Mpla  ORF : Mpla vs. Mten  IRF : Mexa vs. Mten |  | 0.08  0.03  0.04  0.04  0.08  0.07 | 1.40  2.03  1.76  1.08  0.94  3.21 | **0.005**  **0.001**  **0.001**  **0.001**  **0.001**  **0.001** |

**Table S7: Statistical testing for microbial multivariate beta-dispersion of *Millepora* species and habitats.** Results of PERMDISP analyses (nperm = 999) testing for differences in multivariate dispersion across *Millepora* host species and reef habitats. Analyses were performed on CLR-transformed abundances for both Symbiodiniaceae (ITS2) and prokaryotic (16S) assemblages. Pvalues < 0.05 indicate non-homogeneous dispersions between groups.

|  | *Source of variation* | *df* | *F* | *pvalue BETADISPER* |
| --- | --- | --- | --- | --- |
| **Symbiodiniaceae assemblages** | Species | 2 | 0.50 | 0.59 |
|  | Habitats | 2 | 2.35 | 0.11 |
|  | Species x Habitats | 5 | 1.89 | 0.10 |
| **Prokaryotic assemblages** | Species | 2 | 1.48 | 0.23 |
|  | Habitats | 2 | 2.07 | 0.14 |
|  | Species x Habitats | 5 | 3.04 | **0.01** |

**Table S8. Contribution of host genotypes and habitats on Symbiodiniaceae assemblages in *Millepora* species.** Permutational Multivariate Analysis of Variance (PERMANOVA, nperm = 999) testing the effects of host genotypes and reef habitat on CLR-transformed abundances.

| **Symbiodiniaceae assemblages** | *Source of variation* | *df* | *R2* | *F* | *pvalue* |
| --- | --- | --- | --- | --- | --- |
| *M. tenera* | Habitat | 1 | 0.11 | 9.09 | **0.001** |
|  | Genotype (nested) | 8 | 0.46 | 5.26 | **0.001** |
|  | Residuals | 37 | 0.76 |  |  |
| Inner reef flat | *Pairwise comparisons*  G3 vs. G2  G3 vs. G1  G4 vs. G3  G4 vs. G5  G5 vs. G2  G5 vs. G1  G6 vs. G3  G6 vs. G5  G7 vs. G2  G7 vs. G1  G7 vs. G3  G7 vs. G5  Other pairwise comparisons |  |  |  | **0.005**  **0.038**  **0.011**  **0.002**  **0.003**  **0.002**  **0.004**  **0.004**  **0.002**  **0.006**  **0.012**  **0.002**  > 0.05 |
| Outer reef flat | *Pairwise comparisons*  G8 vs. G9 |  |  |  | **0.039** |
| *M.* cf. *platyphylla* | Habitat | 1 | 0.03 | 3.83 | **0.001** |
|  | Genotype (nested) | 9 | 0.54 | 7.99 | **0.001** |
|  | Residuals | 58 | 0.43 |  |  |
| Outer reef flat | *Pairwise comparisons*  G10 vs. G12  G10 vs. G13  G11 vs. G12  G11 vs. G13  G12 vs. G13  G10 vs. G11 |  |  |  | **0.006**  **0.03**  **0.03**  **0.03**  **0.002**  > 0.05 |
| Mid reef slope | *Pairwise comparisons*  G14 vs. G16  G14 vs. G19  G15 vs. G14  G15 vs. G16  G15 vs. G17  G15 vs. G19  G16 vs. G19  G17 vs. G14  G17 vs. G15  G17 vs. G19  G18 vs. G14  G18 vs. G15  G18 vs. G16  G18 vs. G17  G18 vs. G19  G20 vs. G14  G20 vs. G15  G20 vs. G16  G20 vs. G17  G20 vs. G18  Other pairwise comparisons |  |  |  | **< 0.001**  **< 0.001**  **< 0.001**  **< 0.001**  **< 0.001**  **< 0.001**  **< 0.001**  **< 0.001**  **< 0.001**  **0.009**  **< 0.001**  **< 0.001**  **< 0.001**  **0.008**  **0.006**  **< 0.001**  **< 0.001**  **< 0.001**  **0.008**  **0.008**  > 0.05 |

**Table S9. Contribution of host genotypes and habitats on Prokaryotic assemblages in *Millepora* species.** Permutational Multivariate Analysis of Variance (PERMANOVA, nperm = 999) testing the effects of host genotypes and reef habitat on CLR-transformed abundances.

| **Prokaryotic assemblages** | *Source of variation* | *df* | *R2* | *F* | *pvalue* |
| --- | --- | --- | --- | --- | --- |
| *M. tenera* | Habitat | 7 | 0.05 | 2.24 | **0.001** |
|  | Genotype | 1 | 0.20 | 1.27 | **0.001** |
|  | Residuals | 34 | 0.75 |  |  |
| Inner reef flat | *Pairwise comparisons*  G1 vs. G2  G1 vs. G3  G1 vs. G5  G2 vs. G4  G2 vs. G6  G3 vs. G5  G3 vs. G6  G4 vs. G5  G5 vs. G6  G5 vs. G7  G6 vs. G7  Other pairwise comparisons |  |  |  | **0.035**  **0.028**  **0.005**  **0.015**  **0.012**  **0.041**  **0.012**  **0.002**  **0.016**  **0.006**  **0.007**  > 0.05 |
| Outer reef flat | *Pairwise comparisons*  G8 vs. G9 |  |  |  | 0.156 |
| *M.* cf. *platyphylla* | Habitat | 1 | 0.03 | 2.40 | **0.001** |
|  | Genotype | 9 | 0.18 | 1.48 | **0.001** |
|  | Residuals | 58 | 0.80 |  |  |
| Outer reef flat | *Pairwise comparisons*  G10 vs. G12  G10 vs. G13  G11 vs. G12  G11 vs. G13  G12 vs. G13  Other pairwise comparisons  (G10 vs. G11) |  |  |  | **0.006**  **0.023**  **0.008**  **0.035**  **0.008**  > 0.05 |
| Mid reef slope | *Pairwise comparisons*  G14 vs. G15  G14 vs. G16  G14 vs. G17  G14 vs. G19  G15 vs. G16  G15 vs. G17  G15 vs. G18  G15 vs. G19  G16 vs. G17  G16 vs. G18  G16 vs. G19  G16 vs. G20  G17 vs. G18  G17 vs. G19  G17 vs. G20  Other pairwise comparisons |  |  |  | **0.001**  **0.001**  **0.002**  **0.01**  **0.001**  **0.011**  **0.034**  **0.04**  **0.001**  **0.003**  **0.003**  **0.001**  **0.009**  **0.048**  **0.008**  > 0.05 |

**Table S10: Statistical testing for microbial multivariate beta-dispersion in *Millepora* host genotypes and habitats.** Results of PERMDISP analyses (nperm = 999) testing for differences in multivariate dispersion across host genotypes and reef habitats. Analyses were performed on CLR-transformed abundances for both Symbiodiniaceae (ITS2) and prokaryotic (16S) assemblages. Pvalues < 0.05 indicate non-homogeneous dispersions between groups.

| **Symbiodiniaceae assemblages** | *Source of variation* | *df* | *F* | *pvalue BETADISPER* |
| --- | --- | --- | --- | --- |
| ***M. tenera*** | Genotypes | 8 | 3.12 | **0.01** |
|  | Habitats | 1 | 11.68 | **0.003** |
| ***M.* cf. *platyphylla*** | Genotypes | 10 | 2.35 | **0.022** |
|  | Habitats | 1 | 1.14 | 0.27 |
| **Prokaryotic assemblages** |  |  |  |  |
| ***M. tenera*** | Genotypes | 8 | 1.19 | 0.34 |
|  | Habitats | 1 | 3.14 | 0.10 |
| ***M.* cf. *platyphylla*** | Genotypes | 10 | 2.85 | **0.006** |
|  | Habitats | 1 | 6.88 | **0.009** |

**Table S11: Statistical testing for within-genotype co-dispersion**. ANOVA of the linear model testing the influence of within-genotype ITS2 dispersion (Symbiodiniaceae), species and their interaction on within-genotype 16S dispersion (prokaryotes), and likewise the influence of 16S dispersion (prokaryotes), species and their interaction on ITS2 dispersion.

| **Within-genotype co-dispersion** | *Source of variation* | *df* | *SS* | *F* | *pvalue* |
| --- | --- | --- | --- | --- | --- |
| Dispersion 16S | Dispersion ITS2 | 1 | 0.004 | 8E-4 | 0.97 |
|  | Species | 1 | 5.37 | 9.32 | **0.008** |
|  | Disp. ITS2 x Species | 1 | 0.03 | 0.06 | 0.82 |
|  | Residuals | 16 | 9.23 |  |  |
| Dispersion ITS2 | Dispersion 16S | 1 | 0.005 | 8E-4 | 0.99 |
|  | Species | 1 | 4.02 | 6.09 | **0.03** |
|  | Disp. 16S x Species | 1 | 0.04 | 0.06 | 0.81 |
|  | Residuals | 16 | 10.57 |  |  |
